# m5C RNA Methylation Is Dysregulated by Oncogenic Herpesviruses via c-Myc Signaling to Counteract Host Antiviral Factors

**DOI:** 10.1101/2025.05.07.652742

**Authors:** Zhenyu Wu, Dawei Zhou, Zhixian He, Guillaume N. Fiches, Youngmin Park, Jinshan He, Jianwen Chen, Kateepe Arachchige Sudeera Nuwan Shanaka, Thurbu Tshering Lepcha, Prashant Desai, Jian Zhu, Netty G. Santoso

## Abstract

Herpesviruses are a group of double-stranded DNA viruses known to develop versatile viral strategies to escape host immune surveillance for promoting their replication and propagation. This is illustrated by Kaposi’s sarcoma-associated herpesvirus (KSHV), an oncogenic gamma-herpesvirus that overcomes host immune suppression by multiple mechanisms. In this study, we reported that KSHV dysregulates 5-methylcytosine (m5C) modification and mRNA stability of host antiviral factors to benefit its lytic replication. KSHV lytic reactivation or *de novo* challenge led to downregulation of m5C RNA methyltransferases, NSUN2 and NSUN1 (NSUN2/1), while NSUN2/1 depletion promoted KSHV lytic replication. Such KSHV-mediated downregulation of NSUN2/1 is via suppression of the transcriptional factor c-Myc. We further performed the RNA bisulfite sequencing (RNA-BS-seq) to identify KSHV-dependent m5C modification of host mRNAs. KSHV lytic reactivation led to the significant reduction of m5C methylation and mRNA stability of TRIM25, a key activator of the RIG-I pathway, while TRIM25 depletion indeed promoted KSHV lytic replication. These host-virus interaction events were also observed in the infection of another oncogenic gamma-herpesvirus Epstein-Barr virus (EBV). Overall, our results highlighted a new strategy for human gamma-herpesviruses to counteract host antiviral factors and promote their lytic replication by manipulating host m5C RNA methylation.

**Significance Statement:** Our study has identified a novel viral mechanism of human gamma-herpesviruses to manipulate host RNA methylation machineries to subvert immune defenses and enhance viral lytic replication. In particular, our new data showed that KSHV/EBV downregulate the key 5-methylcytosine (m5C) RNA writers NSUN2/1 via c-Myc, and thus decrease m5C modification and stability of TRIM25 mRNA. As TRIM25 is a key E3 ubiquitin ligase in RIG-I signal transduction, its inhibition disrupts RIG-I mediated antiviral sensing and thus favors viral lytic replication. As human gamma-herpesviruses are critical pathogens that highly associate with multiple human diseases especially certain tumors, such studies are significant to shed light in improving the fundamental understanding of virus-host interactions and identifying new host targets for future translational applications.

## Introduction

Human gamma-herpesviruses, including Kaposi’s sarcoma-associated herpesvirus (KSHV) and Epstein-Barr virus (EBV), are well known to evade antiviral immunity, which allows them to establish life-long, persistent infection and cause various tumors^1–4^. KSHV, also known as human herpesvirus-8 (HHV8), is an etiological agent of Kaposi’s sarcoma^5,6^, as well as two lymphoproliferative disorders, primary effusion lymphoma (PEL)^7^ and multicentric Castleman’s disease (CAD)^8^. The life cycle of KSHV has the latent and lytic phases^9^. Once KSHV enters the target cells, primarily B lymphocytes and endothelial cells, it establishes latent infection^10^. At the latent phase, KSHV viral DNA genomes are maintained as episomes, and only a small set of viral genes are expressed^11^. In response to certain stimuli, viral latency can be disrupted and KSHV lytic replication is reactivated. Expression of most KSHV viral genes is induced, and infectious KSHV virions are produced during the lytic phase^12,13^. With the preference to infect epithelial cells and B cells, EBV can also establish latent infection in these cells, which can be reversed to lytic replication^12,14^. Human gamma-herpesviruses develop multiple viral strategies to overcome antiviral immunity, which is well summarized elsewhere^15,16^. To highlight a few examples, KSHV encodes three viral homologues of the interferon-regulatory factor (IRF) family of proteins (vIRF1, 2, and 3), which compete with endogenous IRFs for binding to the promoter regions and block the transcription of type I IFN genes or ISGs^17,18^. KSHV generates viral microRNA miR-K12-7 that reduces the expression of MICB (MHC class I polypeptide-related sequence B) for immune evasion^19,20^. KSHV Open reading frame 10 (ORF10) protein impairs the phosphorylation of STAT1/2 and abolishes the type I IFN signaling^21^. KSHV ORF33 protein hijacks the host cellular phosphatase PPM1G and reduces the phosphorylation of STING and MAVS, thus inhibiting IFN production^22^.

Recently, it has been reported that human gamma-herpesviruses interact with host RNA methylation machineries, whose functions have newly emerged to critically regulate mRNA post-transcriptional events and impact various cellular processes^23,24^. More than 100 types of RNA modifications have been identified so far^25^, while most studies focus on N6-methyladenosine (m6A) methylation. Significant m6A modifications in KSHV viral transcripts were identified^24^, while knockdown of the m6A RNA writer METTL16^26^ or reader YTHDF2^27^ led to enhanced KSHV lytic reactivation. EBV was also shown to downregulate the m6A RNA eraser ALKBH5 to promote its lytic reactivation^28^. We’ve previously identified that another type of RNA methylation, 5-methylcytosine (m5C), plays a role in regulating infection of human immunodeficiency virus (HIV)^29^. We further investigated such RNA modification in the infection of human gamma-herpesviruses. 5-methylcytosine (m5C) is one of the most abundant RNA modifications broadly distributed in diverse RNA species including messenger RNAs (mRNAs), transfer RNAs (tRNAs), ribosomal RNAs (rRNAs), and noncoding RNAs (ncRNAs), as well as viral RNAs^30–33^. The deposition, removal, and recognition of m5C are fulfilled through three groups of proteins, namely writers, erasers, and readers^34^. Writers include the well-defined Nol1/Nop2/SUN domain (NSUN) family of RNA methyltransferases as well as others (members of DNMT and TRDMT family). NSUN2 is identified as the main writer for m5C methylation in mRNAs, while NSUN1 is responsible for m5C methylation primarily in rRNAs^30,35^. Both NUSN2 and NSUN1 (NSUN2/1) were implicated to play a role in controlling viral infections^29,31,36–41^. Erasers (TET family of demethylases)^42^ and readers (YBX1, ALYREF) have also been reported, but their functions in m5C methylation need further verification^43,44^. Recent studies have shown that m5C methylation affects mRNA stability^45^, splicing^46^, nuclear-cytoplasmic export^43^, and translation efficiency^47^, which impact cell proliferation^48^, development^49^, tumorigenesis^44^, as well as antiviral immunity^31,33,41^.

Here we described the previously unappreciated interaction of KSHV infection with host m5C methylation machineries, which led to the downregulation of NSUN2/1 expression through c-Myc transcriptional factor and the reduction of m5C methylation in mRNAs encoding host antiviral factors, such as TIMR25 that is critical to retinoic acid-inducible gene-I (RIG-I) activation^50^. These host-virus interaction events were also observed in EBV infection. Our study identified a new viral strategy for human gamma-herpesviruses to overcome antiviral immunity and promote their replication and spread.

## Results

### KSHV lytic replication reduces NSUN2/1 expression

We initially determined the impact of KSHV lytic reactivation on NSUN2/1 expression. Both the protein and mRNA levels (**Fig. 1A, B**) of NSUN2/1 significantly decreased due to KSHV lytic reactivation from latency in the Doxycycline (Dox) treated TREx.BCBL1.Rta cells, a BCBL1-derived KSHV latency cell line stably transduced with a Dox-inducible RTA expression vector. We ruled out it is due to the effect of Dox, as NSUN2/1 expression remained the same in the Dox-treated BJAB cells, a KSHV-negative B-lymphoma cell line, or the parental BCBL1, a KSHV-positive B-lymphoma line with no Dox-inducible RTA expression (**Fig. S1A**). We also measured NSUN2/1 expression in other KSHV-infected cells, including iSLK.BAC16 and iSLK.r219 cells that contain latently infected KSHV BAC16 and R219 strain respectively, as well as express a DOX-inducible RTA gene^51^. Similarly, NSUN2/1 expression was significantly reduced due to Dox-induced KSHV lytic reactivation in these cells (**Fig.1C, D, sFig. S1B**). Likewise, DOX treatment had no effect on NSUN2/1 expression in the KSHV-negative, parental SLK cell (**Fig. S1C**). We further confirmed these findings in the HEK293.r219 cells transfected with a vector expressing RTA to induce KSHV lytic reactivation (**Fig. 1E, F**). However, RTA gene expression alone without KSHV lytic program failed to cause the decrease of NSUN2/1 expression in either the RTA-transfected parental HEK293 cells (**Fig. S1E**) or the TREx.BJAB.3xFLAG.Rta cells treated with Dox to induce RTA expression (**Fig. S1D**).

**Figure 1.**
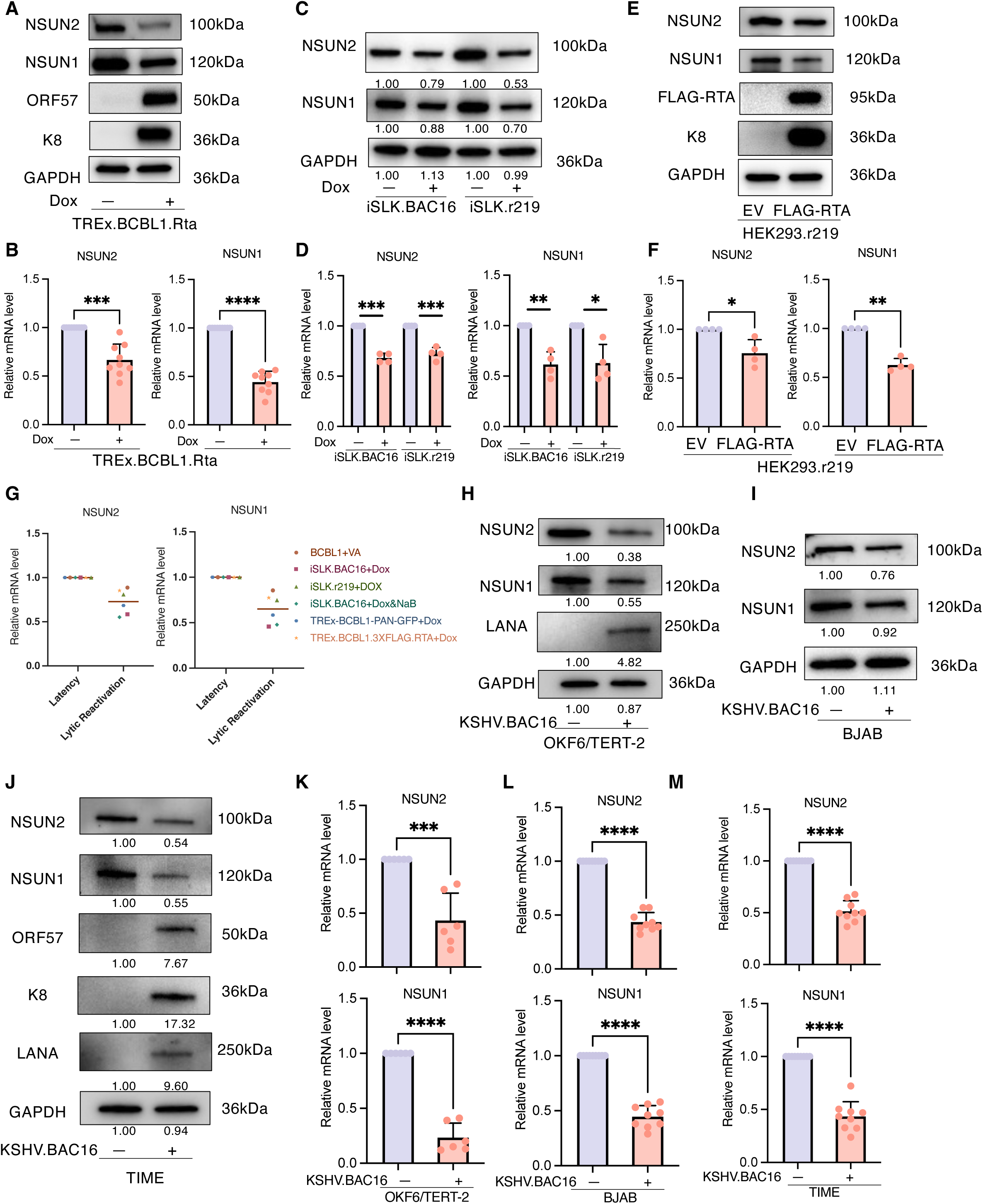
NSUN2/1 expression decreased due to KSHV lytic reactivation and *de novo* infection. (A-C). TREx.BCBL1.Rta cells (A), iSLK.BAC16 and iSLK.r219 cells (B) were treated with Doxycycline or mock. Alternatively, HEK293.r219 cells (C) were transfected with a pcDNA vector expressing FLAG-RTA or empty vector (EV) control. The lysates of above cells were subjected to protein immunoblotting analysis using antibodies recognizing NSUN2, NSUN1,KSHV viral proteins (ORF57, K8), or an anti-FLAG antibody. GAPDH was used as a loading control. (D-F) The RNAs extracted from above cells were subjected to RT-qPCR analysis of NSUN2 and NSUN1, which was normalized to GAPDH. (G) Public-domain RNA-seq data of KSHV-infected cell lines were collected and re-analyzed using the customized pipeline to identify the differentially expressed genes (adjust p-value<0.05 as cutoff). The distinct gene expression level of NSUN2/1 due to KSHV lytic reactivation was illustrated. (H-J) OKF6 cells (H), BJAB cells (I), and TIME cells (J) were inoculated with KSHV.BAC16 viruses. The lysates of above cells were collected at 48 hrs post infection and were subjected to protein immunoblotting analysis using antibodies recognizing NSUN2, NSUN1, and KSHV viral proteins (LANA, ORF57, K8). GAPDH was used as a loading control. Signal intensity of indicated protein bands were quantified using ImageJ. (K-M) The RNAs extracted from above cells were subjected to RT-qPCR analysis of NSUN2 and NSUN1, which was normalized to GAPDH. qPCR results were calculated from three independent experiments and shown as mean ± SD. (\**p* < 0.05, \*\**p* < 0.01, \*\*\**p* < 0.001, \*\*\*\**p* < 0.0001, Student’s t test).

Mining of publicly available RNA-seq datasets generated from KSHV-infected cells further supported our findings (**Table S1**). NSUN2/1 RNA level consistently decreased due to KSHV lytic reactivation in various cell systems from re-analysis of bulk RNA-seq datasets (**Fig. 1G**). With the access to the single-cell (sc) RNA-seq datasets, we also observed that the lower expression of NSUN2/1 associates with KSHV lytic viral transcripts in a small subset of cells (**Fig. S1F-H**). Overall, the above results clearly demonstrated that KSHV lytic reactivation results in the significant reduction of NSUN2/1 expression.

We next determined the impact of KSHV *de novo* infection on NSUN2/1 expression. It is known that multiple cell types are permissive to KSHV *de novo* infection, including epithelial, endothelial cells, and B lymphocytes. Our studies included the OKF6/TERT-2 cells, the immortalized, primary oral mucosal epithelial keratinocytes, as well as the TIME cells, the immortalized, primary endothelial cells. KSHV lytic replication during its initial infection also caused the dramatic decrease of NSUN2/1 protein and mRNA levels in both OKF6/TERT-2 (**Fig. 1H, K, S1I**) and TIME (**Fig. 1J, M, S1J, L**) cells. Such phenotype was also recapitulated in BJAB cells. To increase the infection rate, KSHV.BAC16 virus was concentrated by ultra-centrifugation prior to viral spinoculation with BJAB cells. Consistently, NSUN2/1 expression was downregulated in BJAB cells challenged with KSHV **(Fig. 1I, L, S1K**). Thus, these results indicated that KSHV *de novo* infection also decreases NSUN2/1 expression as another scenario of viral lytic replication.

### Depletion of NSUN2/1 promotes KSHV lytic replication

We speculated that KSHV downregulation of NSUN2/1 may favor its replication. We then measured the impact of NSUN2/1 by its depletion on KSHV lytic reactivation. NSUN2/1 were efficiently knocked down by electroporation of their siRNAs in TREx.BCBL1.Rta cells, which led to the induction of KSHV lytic gene expression, including K8.1, K8, and ORF45 (**Fig. 2G, H**). A similar phenotype was observed in other KSHV-latently infected cell lines, including iSLK.BAC16, iSLK.r219, and HEK293.r219 (**Fig.S2C, D, E**).

**Figure 2.**
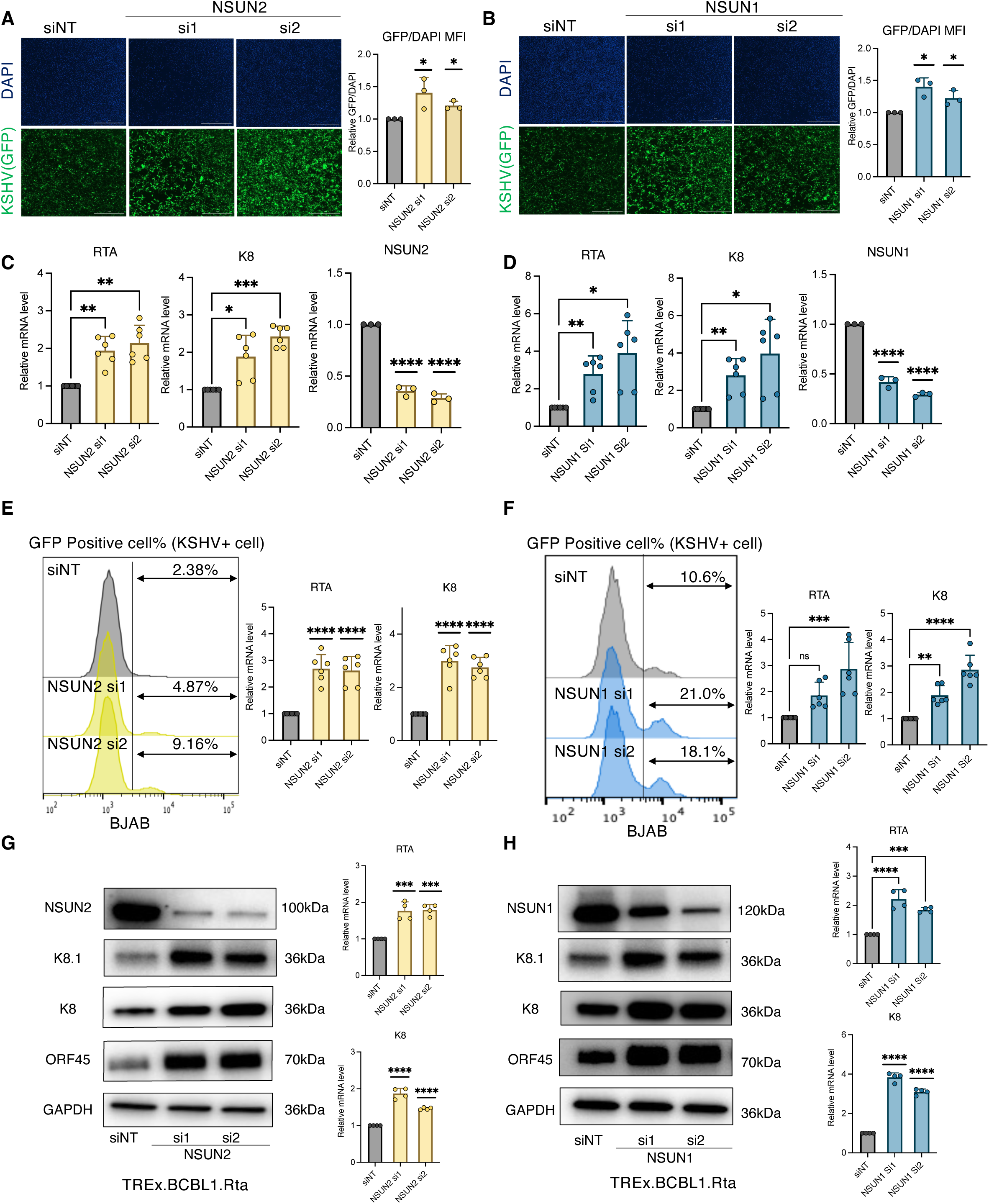
Depletion of NSUN2/1 promoted KSHV *de novo* infection and lytic reactivation. (A-D) TIME cells were transiently transfected with siRNAs (si1, si2) targeting NSUN2 (A) or NSUN1 (B), or non-targeting control siRNA (siNT), followed by inoculation with KSHV.BAC16 viruses. At 48 hrs post infection, these cells were harvested for nuclei staining with Hoechst (blue). GFP fluorescence signal indicating KSHV-infected cells was quantified. (C, D) The RNAs extracted from above cells were subjected to RT-qPCR analysis of NSUN2, NSUN1, and KSHV viral genes (RTA, K8), which was normalized to GAPDH. (E, F) BJAB cells were transiently transfected with siRNAs (si1, si2) targeting NSUN2 (E) or NSUN1 (F), or siNT, followed by inoculation with KSHV.BAC16 viruses. These cells were harvested and subjected to flow cytometry analysis. The percentage of GFP-expressing cells was assessed. The RNAs extracted from above cells were subjected to RT-qPCR analysis of KSHV viral genes (RTA, K8), which was normalized to GAPDH. (G, H) TREx.BCBL1.Rta cells were transiently transfected with siRNAs (si1, si2) targeting NSUN2 (G) or NSUN1 (H), or siNT, followed by protein immunoblotting analysis using antibodies recognizing NSUN2, NSUN1, and KSHV viral proteins (K8, K8.1, and ORF45). GAPDH was used as a loading control. The RNAs extracted from above cells were subjected to RT-qPCR analysis of KSHV viral genes (RTA, K8), which was normalized to GAPDH. qPCR results were calculated from two independent experiments and shown as mean ± SD. (\**p* < 0.05, \*\**p* < 0.01, \*\*\**p* < 0.001, \*\*\*\**p* < 0.0001, Student’s t test).

We also determined whether NSUN2/1 knockdown promoted KSHV *de novo* infection as another scenario. TIME cells were transiently transfected with NSUN2/1 siRNAs, followed by challenging with KSHV.BAC16 viruses (**Fig. 2A, B, C, D, S2A**). Infectivity was measured by quantifying GFP expression from KSHV viral genome or mRNA level of KSHV lytic genes. Our results consistently showed that NSUN2/1 knockdown results in the increase of GFP signals and KSHV lytic gene expression in TIME cells. We observed a similar effect in BJAB cells that were electroporated with NSUN2/1 siRNAs, followed by KSHV.BAC16 spinoculation (**Fig. 2E,F, S2B**). Likewise, to summarize, we demonstrated that depletion of NSUN2/1 favors KSHV lytic replication during both lytic reactivation and de novo infection.

### c-Myc contributes to KSHV-induced downregulation of NSUN2/1

It is intriguing how KSHV turns down NSUN2/1 expression. We reanalyzed the datasets of public RNA annotation and mapping of promoters for analysis of gene expression (RAMPAGE)^52^ that profiles the transcription initiation activity across whole transcriptome^53^. Interestingly, we noticed that promoter activity of NSUN2/1 significantly decreases in KSHV-reactivated iSLK.r219 cells (**Fig. S3C**), indicating that KSHV likely affects NSUN2/1 promoter activity and transcriptional initiation. To gain the insight of NSUN2/1 promoter regulation, we performed the *in silico* analysis of cis-element(s) in the NSUN2/1 promoter region (from -1,000 bp to the transcription start site). By overlapping the results using three public ChIP-seq analytic tools (ChIPBase^54^, ChEA3^55^, hTFtarget^56^), we identified six transcriptional factors (TFs) predicted to associate with NSUN2/1 promoter (**Fig, 3A, B**). Among these TFs, c-Myc is the one with strong and positive correlation with NSUN2/1 expression in the diffuse large B-cell lymphoma (DLBCL) cohort of The Cancer Genome Atlas (TCGA) consortium (r=0.65 for NSUN2, r=0.63 for NSUN1, **Fig. S3A**). Such correlation is also consistent in different tumors and normal tissues (**Fig. S3B**). We reanalyzed the c-Myc CHIP-seq datasets of five lymphoma B cell lines, which confirms the c-Myc binding activity near the NSUN2/1 promoter regions (**Fig. 3C**). Furthermore, the *in silico* scanning by utilizing the motif-based sequence analysis tool FIMO^57^ with the input of c-Myc binding motifs extracted from the open-access database of curated, non-redundant TF binding profiles JASPAR showed that there is indeed a defined c-Myc binding site within the upstream (-1,000bps) to the transcription start site at NSUN2/1 promoters (**Fig. 3D**). We further experimentally demonstrated that c-Myc binding at NSUN2/1 promoters is much lower due to KSHV lytic reactivation by ChIP-PCR assays (**Fig. 3E**).

**Figure 3.**
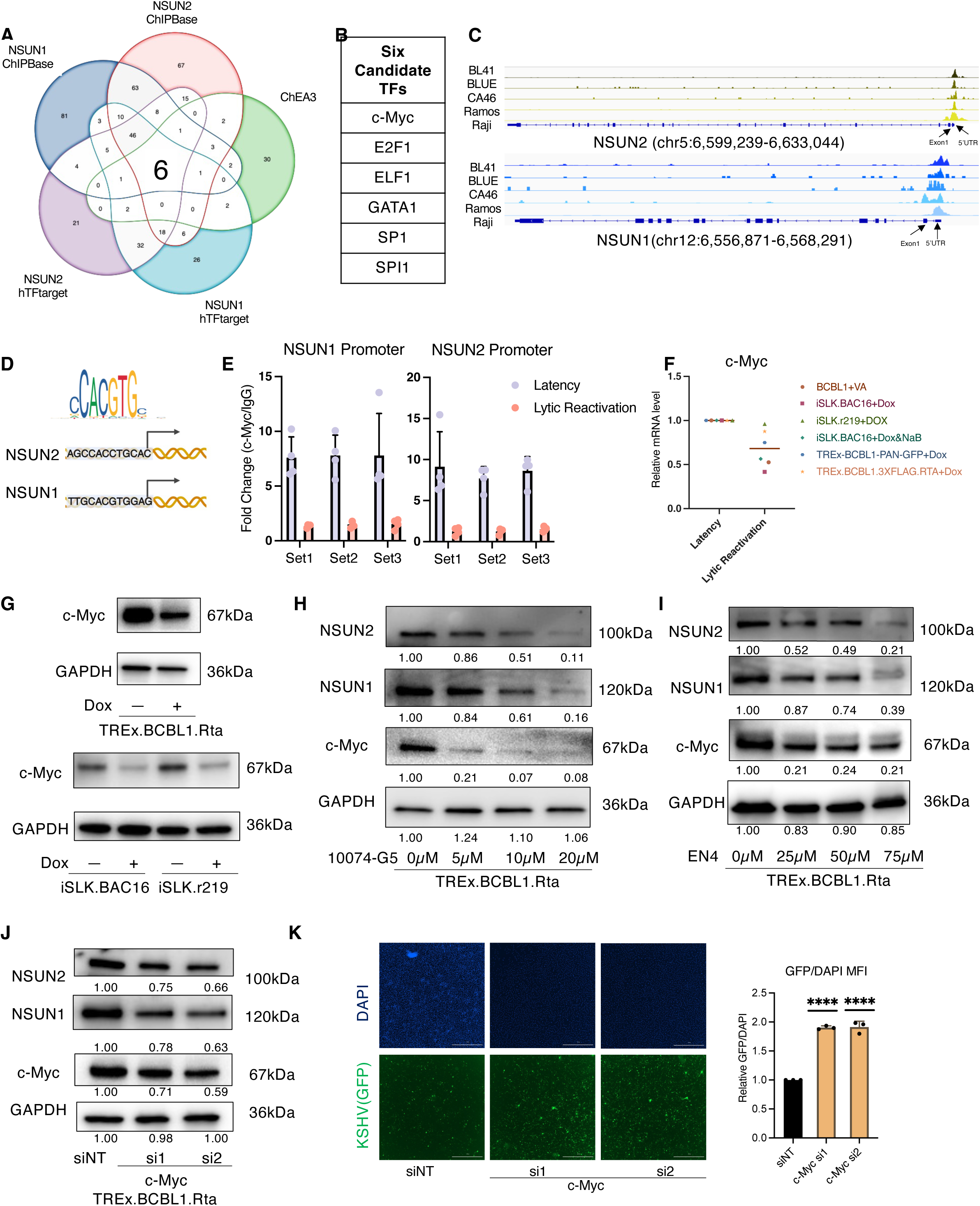
KSHV dysregulated NSUN2/1 expression via c-Myc. (A, B) Transcription factors (TFs) that regulate NSUN2/1 gene expression were predicted by analyzing public-domain ChIP-seq datasets. Venn diagram (A) showed the overlapped TFs across the indicated datasets. Six TF candidates including c-Myc (B) were listed. (C) Integrative genomics viewer (IGV) visualization of c-Myc occupancy near the NSUN2/1 promoter regions using the public-domain c-Myc ChIP-seq dataset of five lymphoma B cell lines (GSE30726). (D) c-Myc binding motif near the promoter regions of NSUN2/1 was illustrated. (E) TREx.BCBL1.Rta cells were subjected to ChIP-PCR analysis for quantification of c-Myc association with the promoter regions of NSUN2/1 using an antibody recognizing c-Myc protein for its immunoprecipitation or a control IgG antibody, followed by qPCR analysis using three sets of primers (Set 1-3) targeting NSUN2 or NSUN1 promoter. (F) Public-domain RNA-seq data of KSHV-infected cell lines were collected and re-analyzed using the customized pipeline to identify the differentially expressed genes (adjust p-value<0.05 as cutoff). The distinct gene expression level of c-Myc due to KSHV lytic reactivation was illustrated. (G) TREx.BCBL1.Rta, iSLK.BAC16, and iSLK.r219 cells were treated with Doxycycline or mock, followed by protein immunoblotting analysis of c-Myc using its specific antibody. GAPDH was used as a loading control. (H, I) TREx.BCBL1.Rta cells were treated with c-Myc inhibitors, EN4 (H) or 10074-G5 (I), at a series of concentrations or mock, followed by protein immunoblotting analysis of c-Myc, NSUN2, NSUN1 using their specific antibodies. (J) TREx.BCBL1.Rta cells were transiently transfected with siRNAs (si1, si2) targeting c-Myc or siNT, followed by protein immunoblotting analysis of c-Myc, NSUN2, NSUN1. (K) TIME cells were transiently transfected with siRNAs (si1, si2) targeting c-Myc or siNT, followed by inoculation with KSHV.BAC16 viruses. These cells were harvested for nuclei staining with Hoechst (blue). GFP fluorescence signal indicating KSHV-infected cells was quantified.

We decided to focus on c-Myc to characterize its role in mediating KSHV-induced NSUN2/1 downregulation. Reanalysis of public RNA-seq datasets revealed that c-Myc expression decreases due to KSHV reactivation (**Fig. 3F**). We verified that c-Myc protein level is significantly lower in KSHV-reactivated TREx.BCBL1.Rta and iSLK cells (**Fig. 3G, S3D).** To further confirm the impact of c-Myc on NSUN2/1 expression in KSHV-infected cells, we transfected c-Myc siRNA in TREx.BCBL1.Rta cells for its knockdown, which led to the obvious reduction of NSUN2/1 protein level (**Fig. 3J**). Alternatively, we utilized two c-Myc inhibitors, EN4^58^, and 10074-G5^59^, to confirm such findings. Both drugs target c-Myc and disrupt its interaction with MAX, thus abolishing c-Myc’s transcriptional function. We observed that these inhibitors effectively decrease both NSUN2/1 and c-Myc protein levels in a dose-dependent manner in TREx.BCBL1.Rta cells (**Fig. 3H, I**), while they caused minimal cytotoxicity (**Fig. S3E, F**). Additionally, similar to NSUN2/1 knockdown the infection rate of KSHV.BAC16 increased in TIME cells transfected with c-Myc siRNA (**Fig. 3K**). Taken all together, we demonstrated that c-Myc expression is reduced due to KSHV lytic replication, which is responsible for KSHV-induced NSUN2/1 downregulation.

### KSHV lytic replication decreases m5C methylation of host mRNAs

Since KSHV lytic replication dramatically downregulates NSUN2/1 that are well known as writers of RNA m5C methylation^43^, we speculated that KSHV may significantly impact m5C methylation of host mRNAs in the infected cells. First, we confirmed that m5C methylation of poly(A)-enriched mRNAs isolated from Dox-treated TREx.BCBL1.Rta cells indeed decreases to nearly 50% by using an anti-m5C antibody in a dot blotting assay (**Fig. 4A**). Next, we performed the RNA bisulfite sequencing (RNA-BS-seq)^43^ studies for the poly(A)-enriched mRNAs isolated from the Dox-treated or un-treated TREx.BCBL1.Rta cells to identify KSHV-dependent landscape of m5C transcriptomic profiles. Bisulfite treatment of RNA samples leads to the conversion of unmethylated cytosines to uracil, which is read as thymine during sequencing, while methylated cytosines (m5C) remain unchanged and read as C during sequencing. RNA-BS-seq is able to detect and map m5C sites at mRNAs at the single nucleotide resolution. For the quality control, C to T conversion rate is a key metric that reflects the efficiency of bisulfite treatment. Our C to T conversion rate was estimated based on that of spike-in mouse Dhfr mRNA, which was all >98% (**Fig. S4A**). Such sequencing analysis identified 2,187 unique m5C sites of host transcripts in un-treated cells, which drastically dropped to 356 sites in Dox-treated, i.e. KSHV-reactivated cells (Fig. 4B). Nealy over 80% m5C sites were lost due to KSHV reactivation. We did not observe any significant alternation regarding the overall distribution of these m5C sites across host genome (**Fig. S4B,C**). These m5C sites were not significantly enriched in specific genomic region or chromosomes, while they would rather occur evenly across all chromosomes. The localization of m5C sites across different regions remains the same with or without KSHV lytic reactivation.

**Figure 4.**
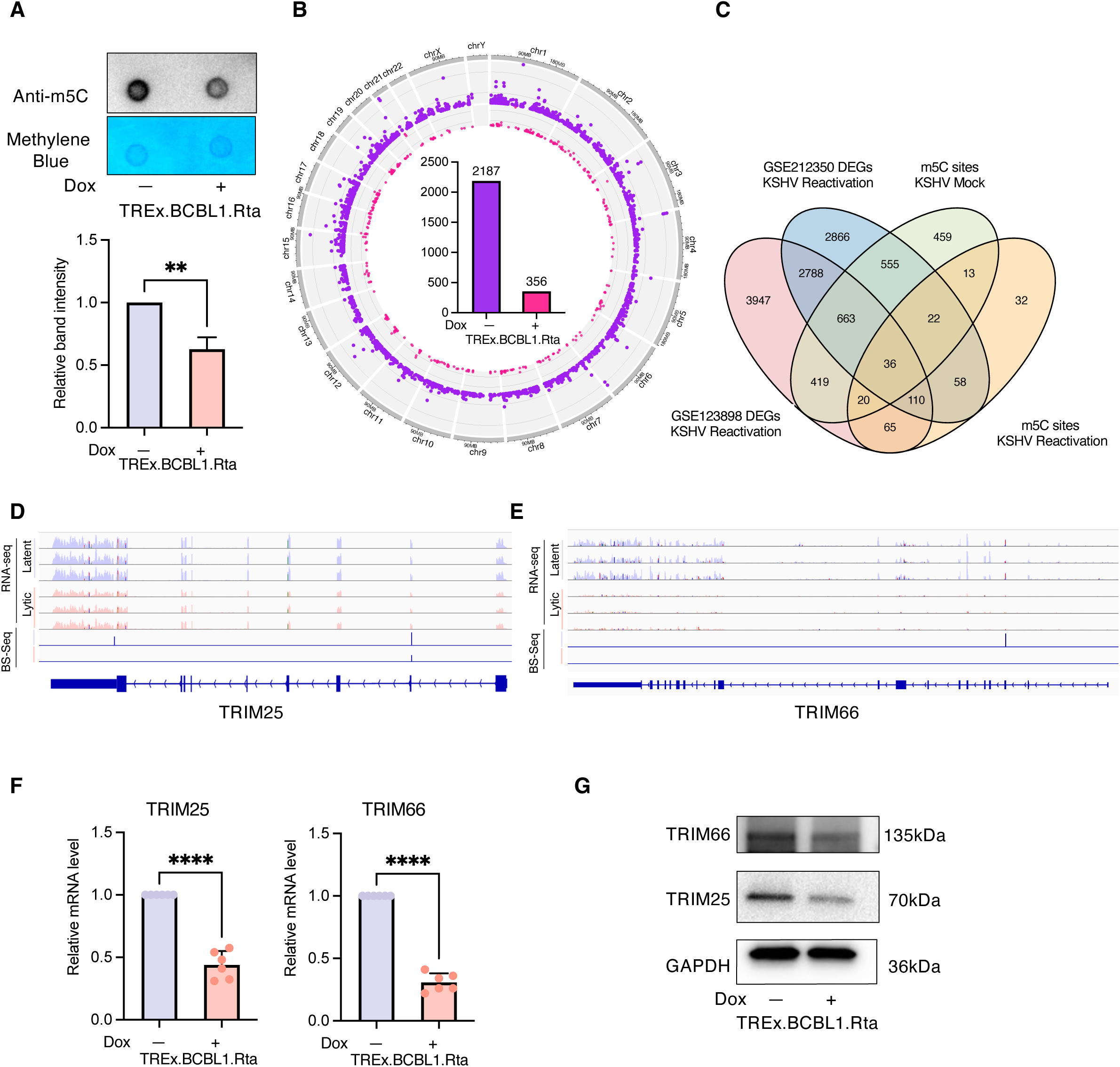
RNA bisulfite sequencing (RNA-BS-seq) analysis identified KSHV-induced dysregulation of m5C modifications at host mRNAs. (A) TREx.BCBL1.Rta cells were treated with Doxycycline or mock. The total RNAs were extracted, followed by mRNA purification. mRNA samples were loaded on positively charged nylon membrane for dot blotting analysis of m5C RNA modification using an anti-m5C antibody or stained with methyl blue to indicate the equal loading. One representative result was shown. The signal intensity of RNA dots was quantified by ImageJ. Results were calculated from three independent experiments and shown as mean +/- SD. (B) Distribution of identified m5C sites at host mRNAs across chromosomes was visualized by Circos plotting. Inner bar charts presented the number of m5C sites with or without KSHV lytic reactivation in TREx.BCBL1.Rta cells. (C) Venn diagram sh owed the overlapped host genes with m5C RNA modification identified from our own RNA-BS-seq studies as well as differentially expressed genes identified from public-domain RNA-seq data of TREx.BCBL1.Rta cells treated with Doxycycline or mock (GSE123898 & GSE212350). (D, E) IGV visualization of identified m5C sites across TRIM25 (D) or TRIM66 (E) mRNAs with the references of RNA-seq data (GSE123898). (F, G) TREx.BCBL1.Rta cells were treated with Doxycycline or mock, followed by RNA extraction and RT-qPCR analysis of TRIM25 and TRIM66 (F), which are normalized to GAPDH. The above cells were subjected to protein immunoblotting analysis (G) of TRIM25 and TRIM66 using their specific antibodies. GAPDH was used as a loading control. (\**p* < 0.05, \*\**p* < 0.01, \*\*\**p* < 0.001, \*\*\*\**p* < 0.0001, Student’s t test).

It has been recently uncovered that RNA m5C modification critically regulates mRNA stability, trafficking, and translation^42–44^. Earlier studies also indicated that m5C methylation is critical to activation of innate immune pathways^60–62^. We then integrated our RNA-BS-seq dataset with those RNA-seq ones generated from TREx.BCBL1.Rta cells with or without induction of KSHV lytic induction, which led to the generation of a list of 36 genes for further investigation (**Fig. 4C**). Pathway analysis revealed that these m5C-hypermethylated transcripts are indeed enriched in the regulation of viral process pathway with a p-value <0.05. Among these hits, we noticed that several genes are from TRIM family, which are significantly subjected to KSHV regulation regarding their mRNA expression and m5C modification (**Table S2**). TRIM43 has already been reported to restrict KSHV infection in an earlier study^63^. However, other TRIM members, like TRIM25, have not been previously reported to impact KSHV infection. Profile of m5C peaks from RNA-BS-seq analysis clearly showed that KSHV lytic reactivation abolishes m5C methylation of their mRNAs (**Fig. 4D,E, S4D,F**). We then measured the mRNA and/or protein levels of TRIM25, TRIM65, TRIM66, which were indeed downregulated due to KSHV lytic reactivation (**Fig. 4F, G, S4E**). Collectively, our sequencing efforts have successfully identified the KSHV-dependent m5C methylation of certain host mRNAs, including a set of TRIM family members (TRIM25, TRIM65, and TRIM66). Additionally, we demonstrated that KSHV lytic reactivation causes no obvious effect on m5C methylation of total RNAs using the dot blotting assays (**Fig. S4G**), indicating that KSHV preferentially affects m5C methylation of mRNAs *vs* other types of RNAs.

### TRIM25 restricts replication of KSHV that reduces its mRNA stability

As TRIM family members (TRIM25, TRIM65, and TRIM66) are targeted by KSHV to downregulate m5C methylation of their mRNAs, we next determined whether these TRIM proteins restrict KSHV lytic replication. Although it is known that TRIM25 is critical to activation of RIG-I signaling that senses the cytosolic viral RNAs, TRIM25 has never been reported to impact KSHV viral replication. Our results showed that knockdown of TRIM25 using its siRNAs efficiently increases the rate of KSHV *de novo* infection in TIME cells by measuring GFP signal (**Fig. 5A**). Similar effect was observed that knockdown of TRIM25 using its siRNAs induces KSHV lytic gene expression in Dox-treated, i.e. KSHV-reactivated TREx.BCBL1.Rta cells (**Fig. 5B, C**). It has been reported that TRIM25 mediates the ubiquitination of RIG-I, thus leading to its activation^50^. We next confirmed that TRIM25 knockdown dramatically reduces the K63-linked ubiquitination of RIG-I but has no effect on its total protein level in TREx.BCBL1.Rta cells (**Fig. 5H**). Knockdown of TRIM25 also induced KSHV lytic reactivation in iSLK.BAC16 and iSLK.r219 cells (**Fig. S5C, E**). Therefore, we identified that TRIM25 is a *bona fide* host restriction factor that limits KSHV lytic replication through modulation of RIG-I ubiquitination and activation. It has already been reported that the knockdown of TRIM65 and TRIM66 promotes KSHV lytic reactivation elsewhere^63^.

**Figure 5.**
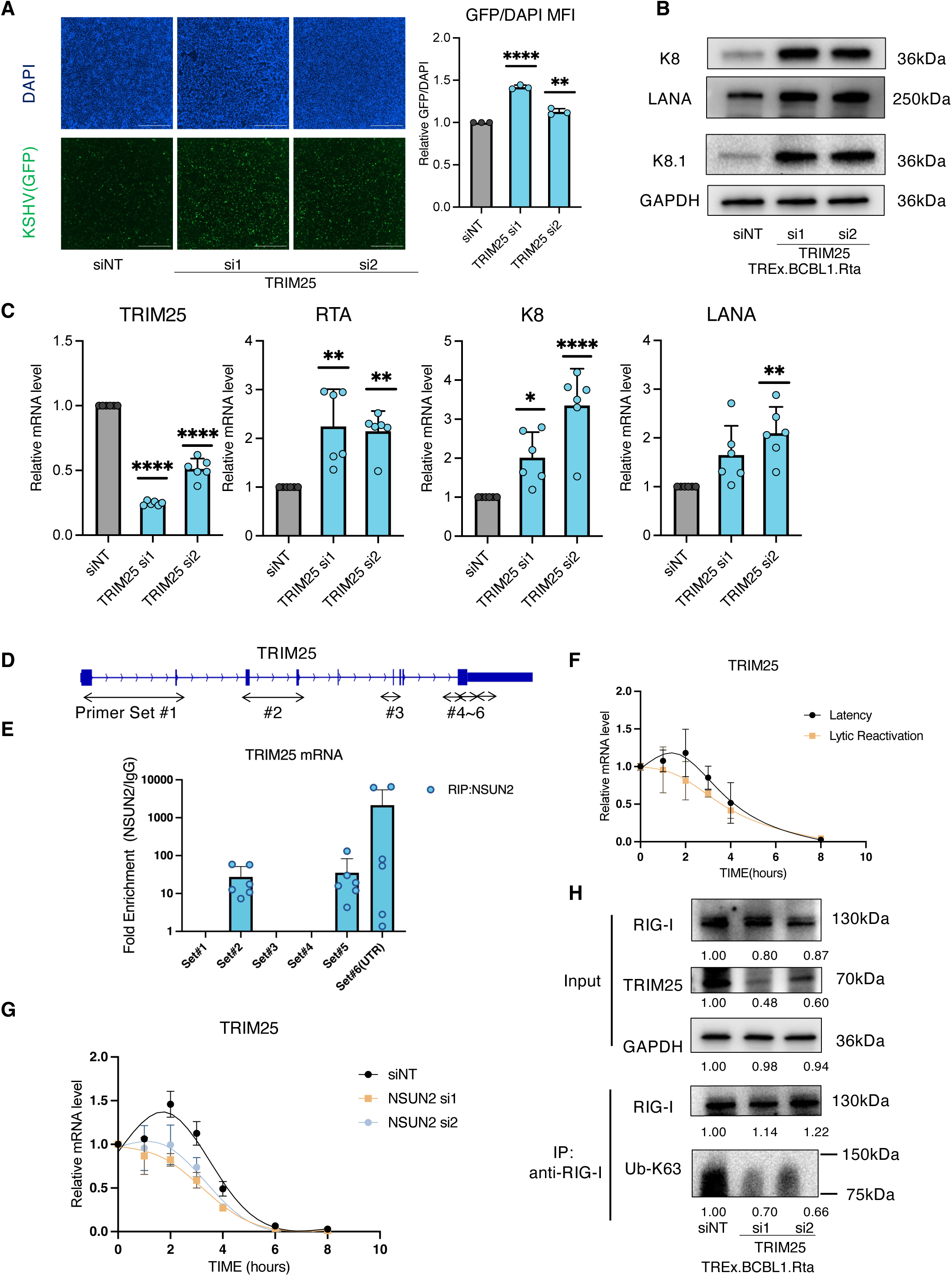
KSHV decreased TRIM25 mRNA stability to antagonize its antiviral activity. (A) TIME cells were transiently transfected siRNAs (si1, si2) targeting TRIM25 or siNT, followed by inoculation with KSHV.BAC16 viruses. At 48 hours post infection, these cells were harvested for nuclei staining with Hoechst (blue). GFP fluorescence signal indicating KSHV-infected cells was quantified. (B, C) TREx.BCBL1.Rta cells were transiently transfected with siRNAs (si1, si2) targeting TRIM25 or siNT, followed by protein immunoblotting analysis (B) using antibodies recognizing KSHV viral proteins (LANA, K8, and K8.1). GAPDH was used as a loading control. The RNAs extracted from above cells were subjected to RT-qPCR analysis (C) of TRIM25 and KSHV viral genes (RTA, K8, and LANA), which was normalized to GAPDH. (D, E) Six sets of qPCR primers (Set1-6) covering the adjunct regions of m5C sites at TRIM25 mRNA were designed (D), which were used to determine their association with NSUN2. TREx.BCBL1.Rta cells were treated with Doxycycline or mock, followed by NSUN2 immunoprecipitation using an anti-NSUN2 antibody or a control rabbit IgG antibody. The co-precipitated RNAs were subjected to RT-qPCR analysis of m5C regions at TRIM25 mRNA. (F, G) TREx.BCBL1.Rta cells were either treated with Doxycycline (F) or transiently transfected with siRNAs (si1, si2) targeting NSUN2 (G), followed by Actinomycin D treatment. Cells were collected at the indicated timepoints. Total RNAs were extracted and subjected to RT-qPCR analysis of TRIM25 RNA level. (H) TREx.BCBL1.Rta cells were transiently transfected with siRNAs (si1, si2) targeting NSUN2 or siNT. Cell lysates were incubated with an antibody recognizing RIG-I antibody or a control mouse IgG. Precipitated protein samples as well as free lysates were analyzed by protein immunoblotting using antibodies recognizing RIG-I, TRIM25, and. K63-specific polyubiquitin. qPCR results of at least three independent experiments were presented as mean ± SD. (\**p* < 0.05, \*\**p* < 0.01, \*\*\**p* < 0.001, \*\*\*\**p* < 0.0001, Student’s t test).

Since we confirmed that TRIM25 represses KSHV lytic replication, we further explored the molecular insights for KSHV to downregulate TRIM25. We first employed the RNA immunoprecipitation (RIP) and qPCR (RIP-qPCR) assays to map the region(s) of TRIM25 mRNAs that associate with NSUN2 protein (**Fig. 5D**). Interestingly, we observed that NSUN2 preferentially binds to two regions near m5C methylation sites of TRIM25 mRNAs, one between exon 3 and 4 and the other at 3’UTR (**Fig. 5E**). We then determined whether mRNA stability of TRIM25 is affected by KSHV downregulation of its m5C methylation. We found that TRIM25 mRNA degrades much faster in TREx.BCBL1.Rta cells at the condition of KSHV reactivation by Dox induction (**Fig. 5F**) or NSUN2 knockdown by its siRNAs (**Fig. 5G**). Knockdown of NSUN2 also reduced TRIM25 expression in iSLK.BAC16 and iSLK.r219 cells (**Fig. S5B, D**). Overall, we demonstrated that KSHV downregulates NSUN2-mediated m5C methylation of TRIM25 mRNA, thus reducing its stability, which would benefit its lytic replication.

### EBV also downregulates NSUN2/1 and counteracts TRIM25’s restriction

To determine whether downregulation of NSUN2/1 expression as well as m5C methylation of TRIM25 mRNA are conserved events, our studies also included another human gamma herpesvirus, EBV. We showed that BJAB cells challenged with EBV.BX viruses have much lower expression of NSUN2/1 (**Fig. 6C, S6A**). The similar phenotype was also observed in human IgG (hIgG) or 12-O-Tetradecanoylphorbol 13-acetate and Sodium Butyrate (TPA+NaB) treated Akata BX cells, a B-lymphoma cell line harboring latently infected EBV.BX viruses, which led to the induction of EBV lytic reactivation (**Fig. 6A, S6C, D**). Reanalysis of several public RNA-seq datasets also supported that EBV lytic reactivation reduces c-Myc and NSUN2/1 expression in Akata BX cells (**Fig. 6B**). On the other hand, we verified that knockdown of NSUN2/1 by their siRNAs increase transcripts of EBV lytic genes (BZLF1, BNRF1) in Akata cells (**Fig. 6D**). Likewise, knockdown of TRIM25 by its siRNAs also induced EBV lytic gene expression in Akata cells (**Fig. 6E**). Therefore, we concluded that EBV also downregulates NSUN2/1 and counteracts TRIM25’s restriction.

**Figure 6.**
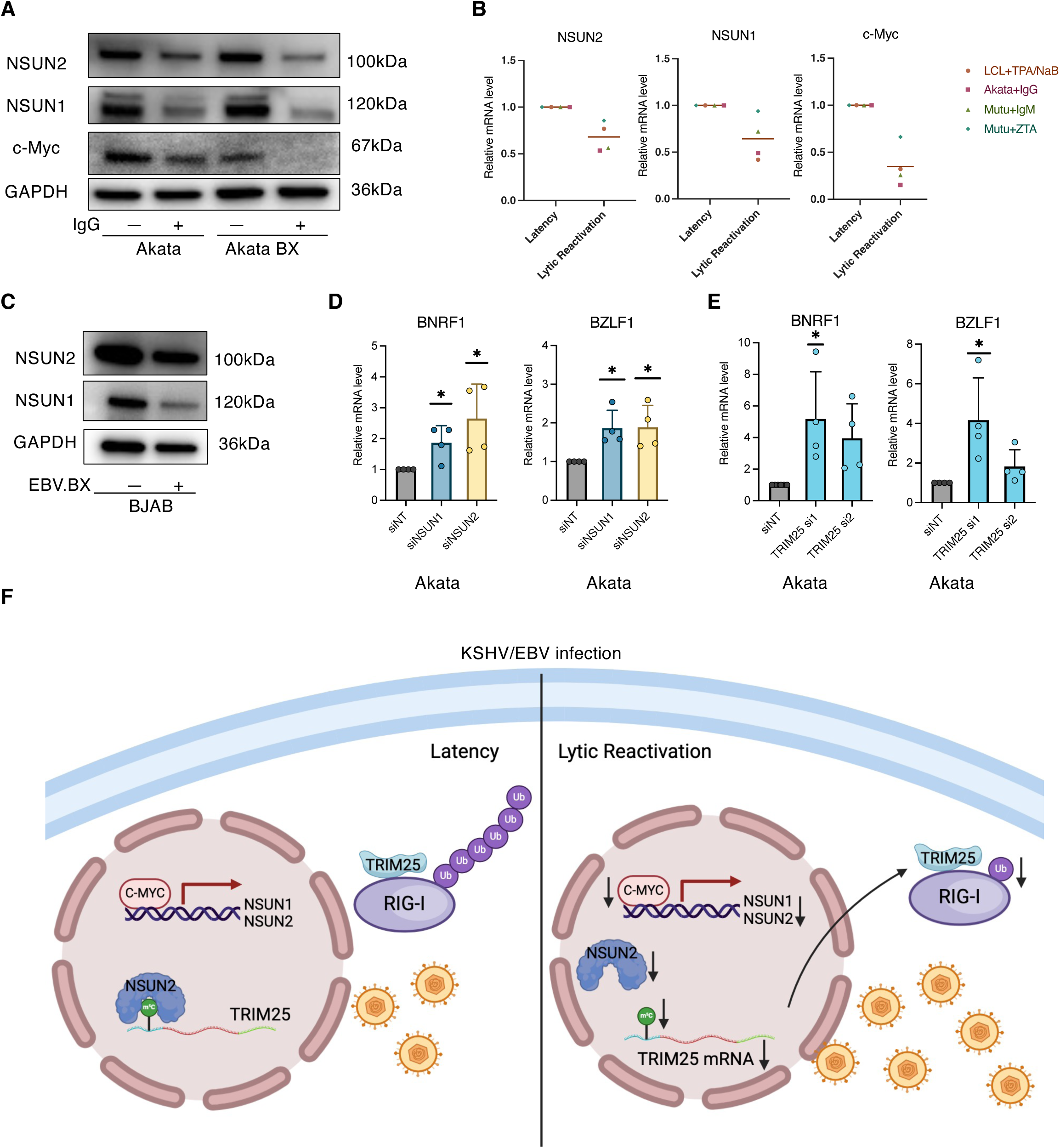
EBV also reduced expression of NSUN2/1 that restrict its lytic infection. (A) Akata and Akata BX cells were treated with a human IgG antibody to induce EBV lytic reactivation, followed by protein immunoblotting analysis using specific antibodies recognizing NSUN2, NSUN1, and c-Myc. GAPDH was used as a loading control. (B) Public-domain RNA-seq data of EBV-infected cell lines were collected and re-analyzed using the customized pipeline to identify the differentially expressed genes (adjust p-value<0.05 as cutoff). The distinct gene expression level of NSUN2/1 due to EBV lytic reactivation was illustrated. (C) BJAB cells were inoculated with EBV.BX viruses. Cell lysates were collected at 48hrs post infection and were followed by protein immunoblotting analysis using specific antibodies recognizing NSUN2 and NSUN1. (D, E) Akata cells were transiently transfected with siRNAs targeting NSUN2/1 (D) or TRIM25 (E), or siNT. The RNAs were extracted and subjected to RT-qPCR analysis of EBV viral genes (BZLF1, BMRF1), which are normalized to GAPDH. (F) A model that human gamma-herpesviruses (KSHV, EBV) downregulate NSUN2/1 and decrease m5C modification of TRIM25 mRNA to favor their lytic replication. c-Myc is downregulated due to KSHV/EBV lytic replication, which reduces NSUN2/1 expression. Coordinately, NSUN2/1-mediated m5C modification of TRIM25 mRNA is inhibited, which impairs its stability. As TRIM25 is a key E3 ubiquitin ligase in RIG-I signal transduction, its inhibition disrupts RIG-I mediated antiviral sensing and thus favors KSHV/EBV lytic replication.

## Discussion

In this report, we identified that expression of m5C RNA writers NSUN2/NSUN1 is downregulated in both scenarios of KSHV lytic reactivation and de novo infection (**Fig. 1**). Reanalysis of RNA-seq datasets from other studies also revealed the similar results. This phenotype occurred in multiple cell types, including B cells, endothelial and epithelial cells, which was also repeatable regardless of approaches to reactivate KSHV, including Dox inducible system^64^ (**Fig. 1A-D**), over-expression of exogenous RTA, and treatment with valproic acid^65^ or sodium butyrate^66^. However, KSHV lytic protein RTA alone was not sufficient to downregulate NSUN2/1 (**Fig. S1D, E**). Interestingly, KSHV lytic reactivation appeared to have no consistent effect on downregulating expression of other NSUN family members (NSUN3-6), either from the re-analysis of public-domain RNA-seq datasets of KSHV latency *vs* lytic reactivation (**Table S5**), or experimentally by RT-qPCR assays of their gene expression in Dox-treated TREx.BCBL1.Rta, iSLK.BAC16, and iSLK.r219 cell models (**Fig. S9A,B**). Furthermore, depletion of other NSUN family members (NSUN3-6) by their siRNAs also failed to cause consistent effect on KSHV viral gene expression in TREx.BCBL1.Rta cells (**Fig. S9C).**

Our results also supported the model that KSHV downregulates NSUN2/1 to favor its lytic replication, as we showed that depletion of NSUN2/1 indeed promotes both KSHV lytic reactivation and *de novo* infection in multiple cell systems (**Fig. 2**). Consistently, depletion of NSUN2/1 enhanced KSHV viral gene expression in TIME cells challenged with KSHV in a time course infection assay (Fig. S7A,B). NSUN2/1 depletion indeed increased the newly synthesized KSHV virions from Dox-treated TREx.BCBL1.Rta cells in a viral titration assay as well (Fig. S7C,D). Interestingly, a related study reported that depletion of NSUN2 and loss of m5C modification resulted in an increased level of EBV viral microRNA EBER1^36^. NSUN2/1 have been implicated to control infection of other viruses as well. We previously showed that NSUN1 restricts HIV-1 replication, suppresses HIV-1 transcription, and promotes viral latency via m5C methylation of viral TAR RNA^29^. HBV also downregulated the expression of NSUN2, while knockdown of NSUN2 significantly decreased virus-induced IFN production and thus enhanced HBV viral replication^37^. Furthermore, Wang et al. reported that NSUN2 specifically methylates four m5C sites at IRF3 mRNA and enhances its expression and stability, thus promoting interferon responses against infection of multiple viruses, including SeV, VSV, HSV, and ZIKV^41^. On the contrary, Courtney et al. reported that NSUN2-mediated m5C RNA modification is critical to retroviral replication^38,40^, although Eckwahl et al. described that knockdown NSUN2 has no effect on HIV viral protein production or release^39^. A recent study revealed that NSUN2 depletion upregulates ncRNAs RPPH1 and 7SL RNAs that are direct ligands for the RIG-I mediated IFN response and thus inhibits the replication of multiple viruses, such as VSV, RSV, hMPV, SeV, and HSV^31^. Overall, these findings indicate that the m5C RNA writers NSUN2/1 may play a profound role in regulating viral infection and antiviral immunity by targeting host and viral mRNAs, which would generate a distinct impact in the milieu of unique host cell types for individual viral species.

To gain more mechanistic understanding how KSHV dysregulates NSUN2/1, our further exploration resulted in a key finding that c-Myc is the host TF that mediates KSHV’s such impact (**Fig. 3**). Through mining of existing RAMPAGE databases as well as *in silico* analysis of chromatin association of TFs and their motif recognition, we pinpointed c-Myc as the key TF that associates with promoters of NSUN2/1 and regulates their transcriptional initiation. We experimentally verified this finding by c-Myc knockdown or use of its inhibitors, which both led to the decrease of NSUN2/1 expression. We also confirmed that c-Myc expression reduces due to KSHV lytic replication, which correlates with KSHV’s impact on NSUN2/1, and that depletion of c-Myc indeed facilitates KSHV lytic gene expression (Fig. S10H,I).

Other studies support our finding that c-Myc is downregulated by KSHV lytic replication. c-Myc is required for the maintenance of KSHV latency, and its knockdown increases RTA promoter activity and promotes KSHV lytic reactivation^67^. During KSHV latency, c-Myc protein is stabilized and its function is stimulated by viral latent proteins, LANA, vIRF3, and vFLIP^68–71^. However, KSHV lytic protein vIRF4 strongly suppresses c-Myc expression during lytic replication ^72^. Thus, c-Myc expression and/or function is overall suppressed by KSHV lytic program. Beyond c-Myc, our analysis also indicated that other TFs may regulate NSUN2/1 expression, although they seemed to show weak or negative correlations (Fig. S3A). For example, E2F1 was identified as a host TF that positively regulates NSUN2 expression^73^. c-Myc is likely the main TF that regulates NSUN2/1 expression, as we showed that KSHV lytic infection has no drastic effect on expression of other three tested TFs, including GATA1, ELF1, and E2F1 (Fig. S10A,B), and that depletion of these TFs does not affect NSUN2/1 expression (Fig. S10C-F) or KSHV viral gene expression (Fig. S10G). However, it remains to be further examined whether other TFs beyond c-Myc also play a role in regulating NSUN2/1 expression.

In contrast to KSHV lytic reactivation, we noticed that c-Myc level remains unchanged while NSUN2/1 expression reduce during KSHV *de novo* infection. We speculated that there might be other transcriptional/epigenetic mechanisms that govern NSUN2/1 expression beyond c-Myc, supported by our findings that multiple TFs are implicated to regulate NSUN2/1 (**Fig. 3A,B**). Other epigenetic factors might also play a role. It has been reported that NSUN1 is regulated by the long noncoding RNA LINC00963 and miRNA miR-542-3p^74^. KDM5A contributed to NSUN2 expression through modulation of H3K4me3 as well^75^. These findings suggest that NSUN2/1 expression could be subjected to a complex regulatory network that involves multiple transcriptional/epigenetic mechanisms, which might vary depending on the cellular environment and/or viral infection status.

Since KSHV dramatically downregulates expression of NSUN2/1 that are the key RNA m5C writers, we expect that it would further substantially affect m5C epitranscriptome in KSHV-infected cells. RNA m5C methylation may be affected by not only writers but also readers and erasers. However, functions of recently identified m5C readers and erasers are still under debate and not widely accepted and/or confirmed yet. To profile m5C epitranscriptome, a set of sequencing approaches can be used, including RNA-BS-seq, 5-azacytidine-mediated RNA immunoprecipitation sequencing (Aza-IP-seq), nanopore sequencing, methylation-iCLIP sequencing (miCLIP-seq), and m5C antibody-based immunoprecipitation sequencing (m5C-RIP-seq)^76–79^. However, we preferred RNA-BS-seq as it is well established, which allows the specific m5C methylation sites to be mapped. To allow the identification of m5C sites that exclusively occur to mRNAs, we purified mRNAs from total RNAs by using polyA selection, as ribosomal RNAs (rRNAs) comprise up to 90% of total RNAs. We also used the mouse Dhfr mRNA as a spike-in positive control to estimate the BS conversion rate of our samples, which was highly efficient with most over 99% (Fig. S4A). We were able to identify nearly three thousand sites that were differentially m5C methylated due to KSHV lytic reactivation. Most of these identified m5C sites existed at the gene body (Fig. S4B). More than 70% of these m5C sites were annotated to mRNAs, while the other 20% to lncRNAs (Fig. S4C). Similar distributions were observed in other RNA-BS-seq studies^80^ . There was no significant difference regarding the distribution pattern of m5C sites across host genomes at the condition of KSHV latency *vs* lytic reactivation in TREx.BCBL1.Rta cells (Fig. S4B, C). Previous studies suggested that the m5C modification is abundant in viral RNAs for certain viruses^38,47,81^. However, we only identified the limited m5C sites in KSHV viral transcripts during its latency and lytic reactivation.

We generated a solid list of host factor candidates for the next-step studies of their role in KSHV infection. As RNA m5C methylation has been reported to affect RNA stability and splicing^46^, nuclear-cytoplasmic RNA export^43^, and mRNA translation efficiency^47^, we would like to characterize how m5C RNA methylation exactly impacts these factors. In this report, we prioritized TRIM25 for further investigation, as it is known as an E3 ubiquitin ligase that ubiquitinates the N terminus of RIG-I, one of the pattern recognition receptors that are specific for viral RNA sensing^82,83^. TRIM25-mediated ubiquitination of RIG-I is essential for its antiviral activity, and it was proven that knockdown of TRIM25 increases infection of Sendai virus in HEK293T cells^84^. Additionally, TRIM25 is also required for ZAP’s antiviral function^85–87^. Recent studies suggested that TRIM25 is an RNA-binding protein that may modulate innate immunity^86^. Multiple viruses, including influenza A virus (IAV)^88,89^, human metapneumovirus (HMPV)^90^, severe acute respiratory syndrome coronavirus (SARS-CoV)^91^, and dengue virus^92^, were implicated to hijack TRIM25 for inhibition of RIG-I signaling. However, TRIM25’s role in KSHV infection has never been investigated previously. Our studies demonstrated that TRIM25 also restricts KSHV lytic replication, while m5C methylation of its mRNA is downregulated by KSHV to reduce TRIM25’s mRNA stability in a NSUN2-dependent manner (**Fig. 5**). We believe that these studies identified a novel viral strategy to target mRNA methylation and turnover of host antiviral factors.

From the RNA-BS-seq studies, we indeed identified a few m5C sites at KSHV viral transcripts (Table S4). However, the abundance of KSHV viral transcripts and their m5C modifications at viral latent stage was fairly low. In contrast, we identified one m5C site at KSHV K5 transcript that occurred at both viral latent and lytic stages with decent coverage. We further determined the impact of the m5C modification at 544 site of K5 on its mRNA level (Fig. S8A,B). We generated the vectors expressing either wild-type or mutated K5 (544CtoA) cDNA for cell transfection assays. Our results showed that depletion of NSUN2/1 has no obvious effects on the ratio of wild-type vs mutated K5 mRNA level, indicating that K5 transcript is not subjected to the regulation of m5C writers. We speculated that KSHV dysregulation of NSUN2/1 might preferentially affect host mRNA m5C methylome.

EBV is another human gamma-herpesvirus that shares significant similarities with KSHV regarding viral life cycle and escape from antiviral immunity^93–95^ . We identified that EBV lytic replication also commonly downregulates NSUN2/1 expression as well as subsequent m5C methylation of TRIM25 mRNA, while depletion of NSUN2/1 or downstream TRIM25 consistently promotes EBV replication at both scenarios of its lytic reactivation and *de novo* infection (**Fig. 6A-E**). c-Myc likely plays a similar role in mediating EBV-induced NSUN2/1 downregulation. Earlier studies showed that overexpression of c-Myc suppresses expression of EBV lytic genes ^96^, while its depletion promotes EBV viral lytic cycle^97^. We revealed a potentially novel epi-transcriptomic role of c-Myc in regulating KSHV/EBV viral life cycle through modulation of RNA modification. Overall, our studies illustrated a new viral strategy for human gamma-herpesviruses KSHV/EBV to overcome host cellular restriction by manipulating m5C RNA methylation (**Fig. 6F**). The current chemotherapies for treating KSHV/EBV-positive B-cell lymphomas still have limitations^98^. There is an emerging interest to induce KSHV/EBV viral lytic program in tumor cells to benefit antitumor immunotherapies^99–102^. Our fundamental findings indicate that inhibition of host RNA m5C methylation may benefit such KSHV/EBV viral oncolytic strategies for immune clearance of tumor cells.

## Material and Methods

### Cells and compounds

Cell lines, including BJAB, BL41, BCBL1, TREx.BCBL1.Rta, TREx.BCBL1.3xFLAG.Rta, TREx.BJAB.3xFLAG.Rta^103^, Akata, Akata 4E3, Akata BX, BL41.B95.8, BL41.P3HR1, were cultured in RPMI supplemented with 10% FBS. Cell lines, including SLK, iSLK.BAC16, iSLK.r219, HEK293, HEK293T, HEK293.r219, AGS, AGS.BX, were cultured in DMEM supplemented with 10% FBS. TIME cells^104^ were obtained from Dr. Shou-Jiang Gao (University of Pittsburgh) and maintained in Vascular Cell Basal medium complemented with Endothelial Cell growth kit (ATCC, Cat#PCS-100-030 and PCS-100-041) as previously described (*3*). OKF6/Tert2 cells^105^ were kindly shared by Dr. James G. Rheinwald, and were cultured in keratinocyte serum-free medium (K-sfm) with supplement (Thermo Scientific, CAT#17005042). TREx.BJAB.3xFLAG.Rta^106^ was kindly shared by Dr. Zsolt Toth. TREx.BCBL1.Rta, iSLK.BAC16, iSLK.r219 cells underwent antibiotic selection every other week as described previously^107^. DMSO (Cat#BP231-100) was obtained from Fisher Scientific. c-Myc inhibitors, EN4, and 10074-G5, were purchased from Sellchem. The cytotoxicity of compounds was determined by using the ATP-based CellTiter-Glo Luminescent Cell Viability Assay (Promega, Cat#G7572) following manufacturer’s instructions and analyzed by the Cytation 5 multimode reader (luminescent mode).

### Quantitative PCR (qPCR) and RNA immunoprecipitation with qPCR (RIP-qPCR)

Total RNAs were extracted using the NucleoSpin RNA extraction kit (Macherey-Nagel, Cat#740955.250) following the protocol provided by the manufacturer. RNA samples were reverse transcribed to cDNAs using iScript (BioRad Cat#1708891). Real-time qPCR was performed on a CFX96 instrument (BioRad), by mixing 5 ul (2x) SYBR Green Supermix (BioRad, Cat#1725214), 0.5 ul primer mix, and 4.5 ul cDNA template. Data was analyzed by using the ΔΔCt method with GAPDH as an internal control. RIP assays were performed following the previous publication^108^. In brief, cells were fixed and lysed using NT2 buffer. Cell lysates were incubated with an anti-NSUN2 antibody absorbed to the magnetic beads (Thermo Scientific, Cat#88803) for overnight at 4 °C. The DNA-free DNA Removal Kit (Invitrogen, Cat#AM1906) was used to remove DNAs. Extracted RNA samples were subjected to the above qPCR analysis. All qPCR primers were listed in **Table S3**.

### Protein immunoblotting and dot blot assays

Protein immunoblotting assays were performed as previously described^109^. The following antibodies were used: anti-GAPDH (Santa Cruz Biotechnology, Cat#sc-47724), anti-NSUN1(Santa Cruz Biotechnology, Cat#sc-398884), anti-NSUN2 (Sigma-Aldrich-Aldrich, Cat#ABE1076), anti-HHV8-K8.1A/B (Santa Cruz Biotechnology, Cat#sc-65446), anti-HHV8-ORF50/RTA (Abbiotec, Cat#251345), anti-KSHV-LANA (Advanced Biotechnologies, cat# 13-210-100), anti-KSHV-ORF45 (Santa Cruz Biotechnology, Cat#sc-53883), anti-KSHV-ORF57 (Santa Cruz Biotechnology, Cat#sc-135746), anti-c-Myc (Santa Cruz Biotechnology, Cat#sc-40), anti-c-Myc for ChIP-PCR (Abcam, Cat#ab56), anti-FLAG (Sigma-Aldrich, Cat#F1804-50UG), anti-TRIM25 (Santa Cruz Biotechnology, CAT#sc-166926), anti-TRIM66 (Santa Cruz Biotechnology, CAT#sc-515177), anti-mouse-HRP (Cell Signaling technology, Cat#7076), anti-rabbit-HRP (Cell Signaling technology, Cat#7074), anti-rat-HRP (Invitrogen, Cat#A18739). Dot blot assays were conducted following the published protocol^110^ with slight modifications. In brief, total RNAs were extracted from cell pellets using Trizol (Thermo Scientific), and mRNAs were further extracted from total RNA using mRNA Dynabeads purification kit (Invitrogen, Cat#61006). RNA concentration was measured by NanoDrop. 300ng of mRNA samples were dropped directly onto the Hybond-N+ membrane (Thermo Scientific, Cat#77016), which was further crosslinked to the membrane using a Stratalinker 2400 UV Crosslinker by twice with the mode (2,400 microjoules [x100]; 25-50 sec). The membrane was then washed and blocked with 5% BSA and then incubated with an anti-m5C antibody (1:1000 dilution) in 10 ml of the antibody dilution buffer for overnight at 4 °C with shaking. The images were captured using the same method as protein immunoblotting assays.

### Transfection and electroporation assays

Turbofect (Thermo Scientific, Cat#R0531) reagents were used for plasmid transfection following the manufacturer’s recommendation as described previously^107^. The plasmid expressing Flag-ORF50/RTA was a gift from Dr. Pinghui Feng ^111^ (University of South California). The vectors expressing TRIM25, KSHV K5 (wild-type or 544CtoA mutant) were generated by GenScript. siRNAs targeting NSUN1, NSUN2, c-Myc, TRIM25, TRIM28, TRIM65, and TRIM66 were requested from Invitrogen. To introduce siRNAs in TIME and OKF6/Tert2 cells, reverse transfection was performed using Lipofectamine RNAiMAX (Invitrogen, Cat#13778100) as previously described^112^. To introduce siRNAs in TREx.BCBL1.Rta, BJAB, BL41 cells, electroporation was conducted using the Invitrogen Neon system following the manufacture’s protocol with the customized parameters. In brief, cells were washed with PBS and resuspend in buffer R with a final concentration of 2 × 10^7^ cells/ml. The setting (1,600 V, 10 ms, 3 pulses) was used for TREx.BCBL1.Rta cells, while the setting (1,350 V, 40 ms, 1 pulse) was for BJAB cells. Knockdown efficiency was verified by protein immunoblotting or RT-qPCR assays.

### Viral preparation and infection

KSHV.BAC16 viruses were generated using iSLK.BAC16 cells as previously described^107^ with slight modifications. In brief, iSLK.BAC16 cells were treated with 1µg/ml Dox and 1mM NaB for 24 hours, and then kept in fresh media containing 2 µg/ml Dox. Dox was refreshed every 2 days. Supernatants were collected at Day 3, 5, 7, and 9, centrifuged (500x g) for 10 mins to remove cellular debris, filtered through the 0.45um filter, and stored at -80 °C. For infection of TIME and OKF6/Tert2 cells, these cells were seeded in 12-well plates one day prior to the experiments at the concentration of 0.3 million of cells per well. Cells were then spinoculated (2500 rpm) with KSHV.BAC16 viruses along with 0.8 µg/ml polybrene for 2 hours at 37 °C. Cell media was removed at 1 hour post of infection, and unbound viruses were washed away. For infection of BL41 and BJAB cells, KSHV.BAC16 viruses were concentrated by ultra-centrifugation using a Type 70 Ti Fixed-Angle Rotor and Beckman Optima L-90K ultracentrifuge at the setting of 24,000 rpm, 2 hours at 4 °C. Cells were spinoculated with KSHV.BAC16 viruses. EBV.BX viruses were generated using AGS.BX cells as previously described ^112^ with slight modifications. AGS.BX cells were treated with 12-O-Tetradecanoylphorbol 13-acetate and Sodium Butyrate (TPA 20 ng/ml and NaB 1mM) for 24 hours. Cells were washed and replenished with fresh media for another 24 hours. Supernatants were collected and centrifuged (500x g) for 10 mins to remove cellular debris, filtered through the 0.45µm filter, and stored at -80 °C. For infection of BJAB and BL41 cells, EBV.BX viruses were also concentrated with the similar method as KSHV prior to spinoculation.

### Protein immunofluorescence and flow cytometry

Protein immunofluorescence assays in cells were performed as previously described^109^ using an anti-NSUN2 or anti-NSUN1 primary antibody, followed by the fluorescently conjugated, anti-Mouse Alexa Fluor 488 or anti-Rabbit Alexa Fluor 647 secondary antibody. Cells were then analyzed by flow cytometry using an Accuri C6 Plus (BD Biosciences). Forward Scatter (FS) and Side Scatter (SS) were used for gating of intact, single cells. Analysis was performed using the FlowJo v10 software. Both KSHV.BAC16 and AGS.BX viruses carry a GFP reporter in the viral genome. For cells infected these viruses, fluorescence was quantified using a BioTeK plate reader with the GFP and DAPI channels as previously described^109^, which was analyzed using ImageJ software.

### RNA stability assays

TREx.BCBL1.Rta cells were treated with Dox (2 µg/ml) and Actinomycin D (10 µg/ml) in the fresh media. Cell samples were collected at 0, 1, 2, 3, 4, 8 hr post of drug treatment, and subjected to RNA extraction. RNA samples were analyzed by qPCR assays. The relative level of TRIM25 mRNA was normalized to 16s rRNA. To determine the effect of NSUN2/1 knockdown, Actinomycin D (10µg/mL) was added to TREx.BCBL1.Rta cells at 72 hours post siRNA electroporation.

### RNA-BS-seq studies

We performed RNA-BS-seq studies to map m5C methylation sites of mRNAs isolated from cells with or without KSHV lytic reactivation. Briefly, total RNAs were extracted from cells using Trizol (Thermo Scientific), and mRNAs were further extracted using using mRNA Dynabeads purification kit (Invitrogen) following the manufacture’s protocol. RNA concentration was measured by NanoDrop. DNA-free DNA Removal Kit (Invitrogen, Cat#AM1906) was utilized to avoid DNA contamination. 500 ng of cellular mRNAs and 1 ng *in vitro* transcribed mouse Dhfr mRNA were mixed as input for RNA bisulfite conversion using EZ RNA Methylation Kit (Zymo R5001). The conversion rate was calculated based on Sanger sequencing results of four internal controls (COL4A5, FAM129B, FURIN, PLOD3) and one external control (Dhfr) following the previous publication^43^. The converted mRNAs were then submitted to GENWIZ for library preparation and sequencing. Data analysis was performed following the previous publication ^44^ with slight modifications. Briefly, clean reads were mapped to the hg38 genome using meRanGh from meRanTK^113^. Conversion rate was calculated based on the mouse Dhfr mRNA spike-in, which revealed the C to T conversion rate >98%. The m5C sites were called using meRanCall from meRanTK (false discovery rate < 0.01). Only sites with a coverage depth ≥ 20, methylation level ≥ 0.2, and methylated cytosine depth ≥ 5 were considered credible. The m5C annotation was performed using meRanAnnotate command with the customized genomic sequences consisting of both human and KSHV genomes. For pathway analysis, differentially methylated mRNAs were subjected to the pathway enrichment analysis using the R package clusterProfiler^114^ to calculate the significant canonical pathways (*p* ≤ 0.01). The RNA-BS-seq data generated from these studies were deposited in the Gene Expression Omnibus database under accession number GSE268135. The code used for above analyses was deposited at github: https://github.com/ZhenyuWu-OSU

### Reanalysis of public RNA-seq, scRNA-seq and ChIP-seq datasets

Public RNA-seq datasets (GSE123898, GSE212350, GSE172275, GSE200781, GSE201000, GSE194239) raw reads fastq files were obtained from GEO database and reanalyzed. To analyze bulk RNA-seq datasets, raw reads quality was assessed by fastp^115^. Reads were trimmed with adapters and aligned to the human genome (GRCh38.p14) with HISAT2 aligner. Uniquely aligned reads were submitted as inputs for mapping to gencode.v39 annotation. For identification of differential gene expression, DESeq2 was run with raw read counts obtained from FeatureCounts. Adjusted p value less than 0.05 was used for differentially expressed gene filtering. To analyze single-cell RNA-seq datasets, expression matrix was downloaded from GEO with the accession number GSE12688 and GSE190558. Low-quality cells or empty droplets were filtered out based on the number of unique genes, the total number of molecules, and the percentage of mitochondrial reads detected in each cell. Normalization and scaling of cell clustering and visualization were performed using SeuratV4.0^116^. Cell identity was assigned based on the metadata and marker genes. ChIP-seq dataset (GSE30726) raw data was acquired from GEO and processed with ENCODE ChIP-seq-pipeline2. Peaks were visualized using IGV.

### Statistics

Experiment results were statistically analyzed by Graphpad PRISM9.0 using the unpaired, two-tailed Student’s t test or the one-way analysis of variance (ANOVA). Variables were compared among the outcomes, and the P values below 0.05 were considered significant.

## Acknowledgment

We would like to thank Drs. Prashant Desai, Jae Jung, Shou-Jiang Gao, and Renfeng Li for the gifts of HEK293.r219, TREx BCBL1.Rta, iSLK.BAC16, OKF6/TERT-2, Akata, Akata BX, AGS and AGS.BX cells, respectively. SLK and BCBL-1 cells were acquired through the NIH AIDS Reagent Program. We thank Drs. Jay A. Levy and Sophie Leventon-Kriss for SLK cells, and Drs. Michael McGrath and Don Ganem for BCBL-1 cells. We also sincerely appreciate the support from Dr. Meng Wang (Research Institute at Nationwide Children’s Hospital) to help us establish the analytic pipelines for mining of ChIP-seq data, and Jianwen Que (Columbia University) for helpful comments and feedback. This study was supported by NIH research grants R03DE029716 and R01CA260690 to N.S.; R01MH134402, R01DA059538, R56AI181631, and R56AI157872 to J.Z.

## Author contributions

J.Z. and Z.W. conceived and designed this study; Z.W. performed most of the experiments; D.Z. and Z.H. provided certain experimental supports; Z.W., N.S., and J.Z. analyzed the results; G.F., Y.P., J.H., J.C., K.S., T.L., P.D. contributed reagents and/or provided advises regarding this study; Z.W. and J.Z. wrote the manuscript; N.S. and J.Z. reviewed the manuscript; N.S. and J.Z. supervised the entire study.

## Data and materials availability

All data needed to evaluate the conclusions in the paper are present in the paper and/or the Supplementary Materials. Additional data and materials related to this paper may be requested from the authors.

---

**Figure S1.**
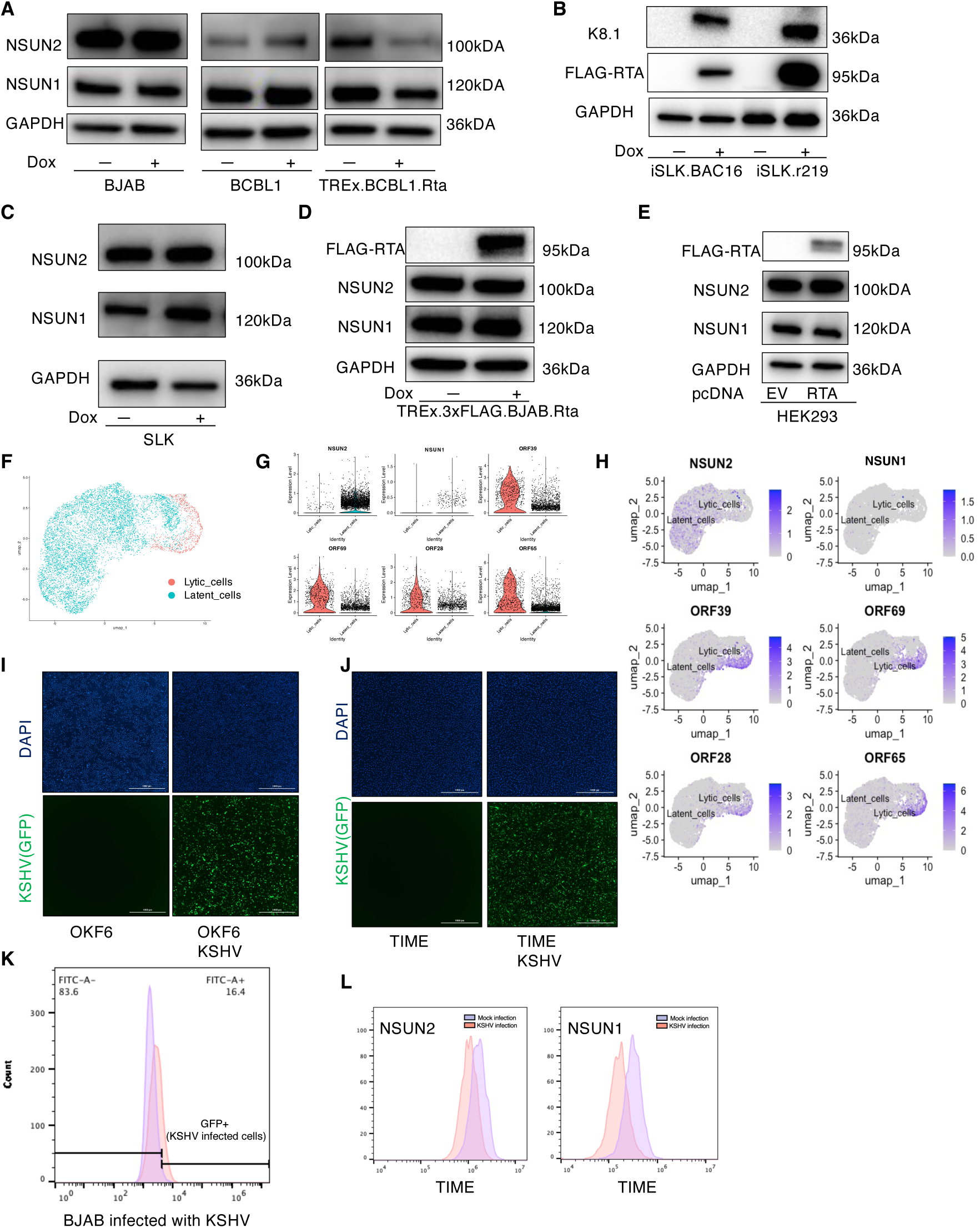
Effect of KSHV lytic reactivation, de novo infection and RTA expression on NSUN2/1. (A) BJAB, BCBL1 and TREx.BCBL1.Rta cells were treated with Doxycycline, followed by protein immunoblotting analysis of NSUN2 and NSUN1. GAPDH was a loading control. (B) iSLK.BAC16 and iSLK.r219 cells were treated with Doxycycline to induce KSHV lytic reactivation, followed by protein immunoblotting analysis of FLAG-RTA and KSHV viral protein K8.1. (C) SLK cells were treated with Doxycycline, followed by protein immunoblotting analysis of NSUN2 and NSUN1. (D) TREx.BJAB.3xFLAG.Rta cells were treated with Doxycycline, followed by protein immunoblotting analysis of NSUN2, NSUN1, and FLAG-RTA. (E) HEK293 cells were transfected with a pcDNA vector expressing FLAG-RTA or empty vector. Protein expression of NSUN2, NSUN1, and FLAG-RTA was measured by protein immunoblotting. (F-H) Reanalysis of public-domain single-cell RNA-seq data of iSLK.r219 cells treated with Doxycycline (GSE190558). Cells with KSHV latent or lytic infection were identified based on the expression of KSHV viral transcripts (F). Expression level and distribution of NSUN2 and NSUN1 as well as indicated KSHV viral lytic genes were plotted using SeuratV4 (G, H). (I, J) OKF6 cells (I) or TIME cells (J) were inoculated with KSHV.BAC16 viruses. These cells were harvested for nuclei staining with Hoechst (blue). GFP fluorescence signal indicating KSHV-infected cells was quantified. (K) BJAB cells were inoculated with KSHV.BAC16 viruses. Percentage of GFP-expressing cells were assessed by flow cytometry. (L) TIME cells were inoculated with KSHV.BAC16 viruses, followed by NSUN2/1 protein immunofluorescence assay and flow cytometry analysis.

**Figure S2.**
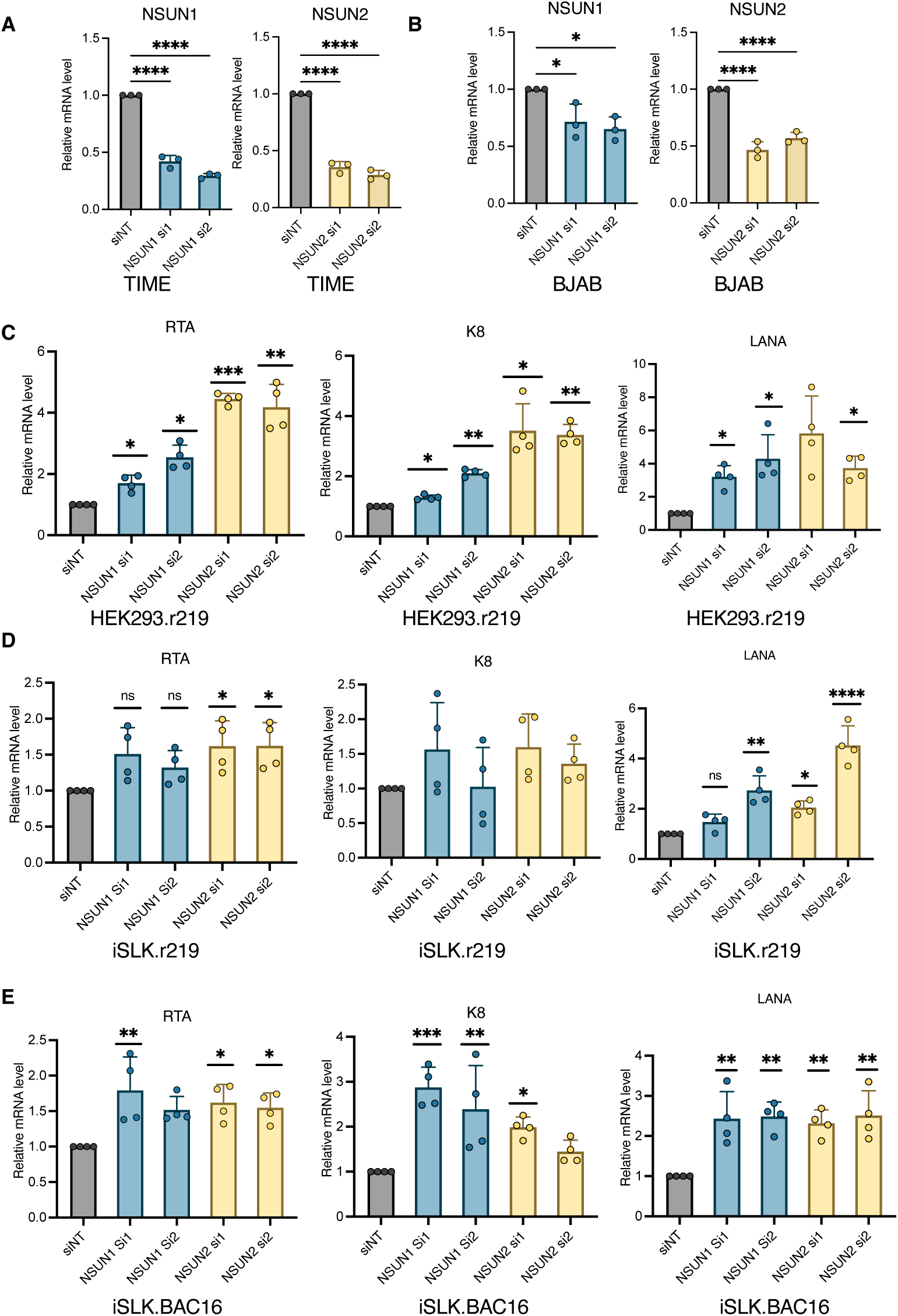
Contribution of NSUN2/1 to the KSHV reactivation. (A, B) TIME (A) and BJAB (B) cells were transiently transfected with siRNAs (si1, si2) targeting NSUN2/1 or siNT. RNAs were extracted and subjected to RT-qPCR analysis of NSUN2/1, which was normalized to GAPDH. (C-E) HEK293.r219 (C), iSLK.r219 (D), and iSLK.BAC16 (E) cells were transiently transfected with siRNAs (si1, si2) targeting NSUN2/1 or siNT. RNAs were extracted and subjected to RT-qPCR analysis of KSHV viral genes (RTA, K8, LANA), which was normalized to GAPDH. qPCR results from at least two independent experiments were presented as mean ± SD. (\**p* < 0.05, \*\**p* < 0.01, \*\*\**p* < 0.001, \*\*\*\**p* < 0.0001, Student’s t test).

**Figure S3.**
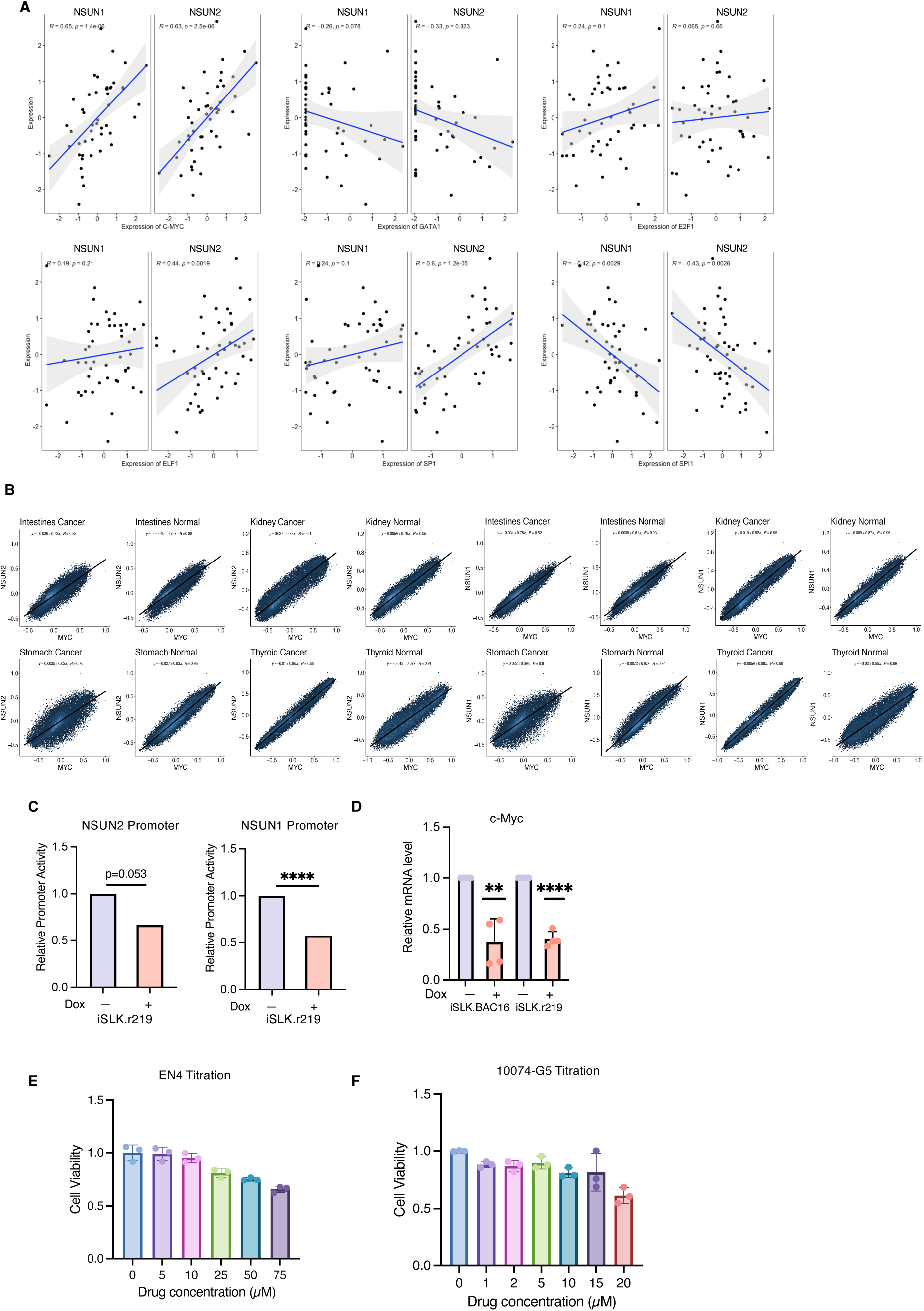
(A) The expression of c-Myc *vs* NSUN2/1 from DLBCL patient cohort of TCGA consortium was plotted, and the Spearman’s correlation of their expression was accessed using R. (B) The expression of c-Myc *vs* NSUN2/1 in multiple cancers and normal tissues were plotted using Correlation AnalyzeR. (C) The promoter activity of NSUN2/1 at the condition of KSHV lytic reactivation *vs* latency was assessed by reanalysis of public-domain RAMPAGE dataset (GSE129902). (D) iSLK.Bac16 and iSLK.r219 cells were treated with Doxycycline or mock. RNAs were extracted and subjected to RT-qPCR analysis of c-Myc, which was normalized to GAPDH. (E, F) Cell viability of TREx.BCBL1.Rta cells were treated with EN4 (E) or 10074-G5 (F) with a series of concentrations, followed by the CellTiter-Glo® luminescent cell viability assay. qPCR results from at least two independent experiments were presented as mean ± SD. (\**p* < 0.05, \*\**p* < 0.01, \*\*\**p* < 0.001, \*\*\*\**p* < 0.0001, Student’s t test).

**Figure S4.**
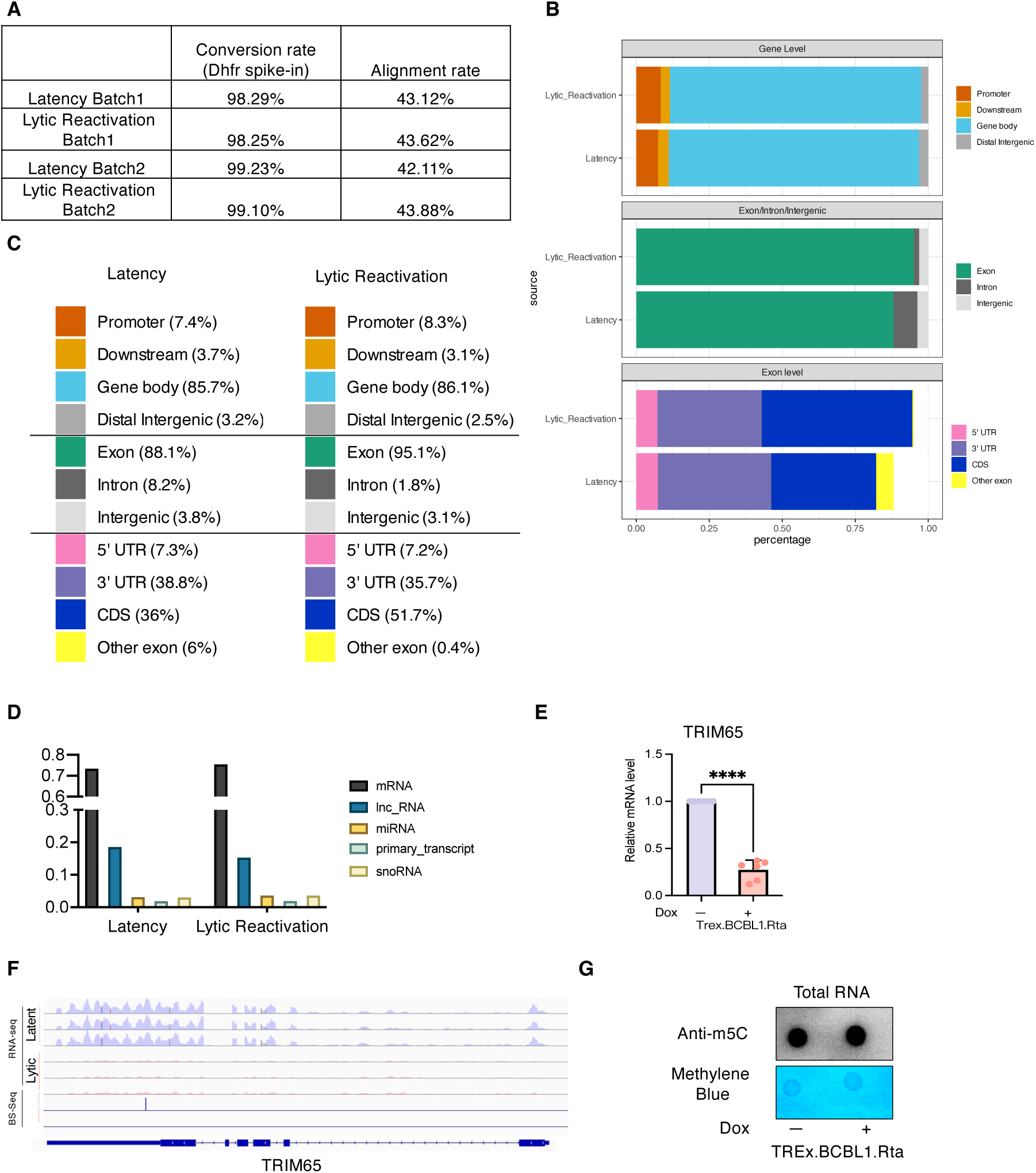
(A) The conversion rate of bisulfite-treated RNA samples were accessed and calculated based on that of the spiked-in mouse Dhfr mRNA prior to submission for RNA-Bis-seq analysis. (B, C) Localization of m5C sites (B) and their percentage (C) at host mRNAs in TREx.BCBL1.Rta cells with or without KSHV lytic reactivation were illustrated using the R package ChIPpeakanno. (D) Distribution of m5C sites across the indicated RNA species in TREx.BCBL1.Rta cells with or without KSHV lytic reactivation were shown as bar charts. (E) Expression of TRIM65 in Doxycycline or mock-treated TREx.BCBL1.Rta cells were measured using RT-qPCR analysis. (F) IGV visualization of identified m5C sites across TRIM65 mRNA with the reference of RNA-seq data (GSE123898). (G) TREx.BCBL1.Rta cells were treated with Doxycycline or mock. The total RNAs were extracted and then loaded on positively charged nylon membrane for dot blotting analysis of m5C RNA modification using an anti-m5C antibody or stained with methyl blue to indicate the equal loading. One representative result was shown. qPCR results from at least three independent experiments were presented as mean ± SD. (\**p* < 0.05, \*\**p* < 0.01, \*\*\**p* < 0.001, \*\*\*\**p* < 0.0001, Student’s t test).

**Figure S5.**
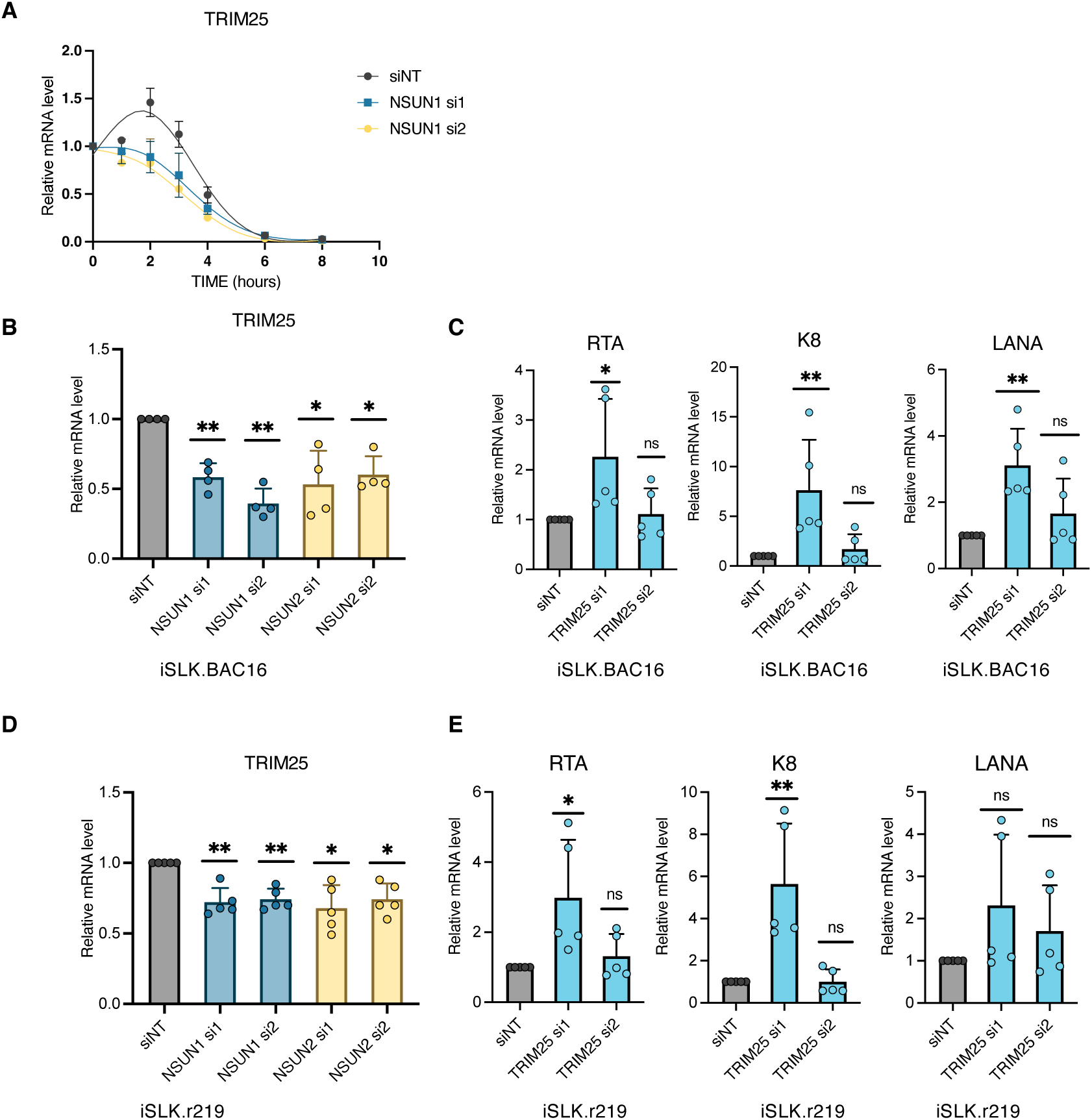
(A) TREx.BCBL1.Rta cells were transiently transfected with siRNAs (si1, si2) targeting NSUN1 or siNT, followed by Actinomycin D treatment. Cell samples were collected at the indicated time points and subjected to RNA extraction and RT-qPCR analysis of TRIM25 mRNA level, which was normalized to GAPDH. (B,C). iSLK.BAC16 cells were transiently transfected with siRNAs targeting NSUN2/1 (B) or TRIM25 (C), followed by RNA extraction and RT-qPCR analysis of NSUN2/1 (B) or indicated KSHV viral genes (C), which was normalized to GAPDH. (D, E). The similar studies were performed for iSLK.r219 cells as (B,C). qPCR results from at least two independent experiments were presented as mean ± SD. (\**p* < 0.05, \*\**p* < 0.01, \*\*\**p* < 0.001, \*\*\*\**p* < 0.0001, Student’s t test).

**Figure S6.**
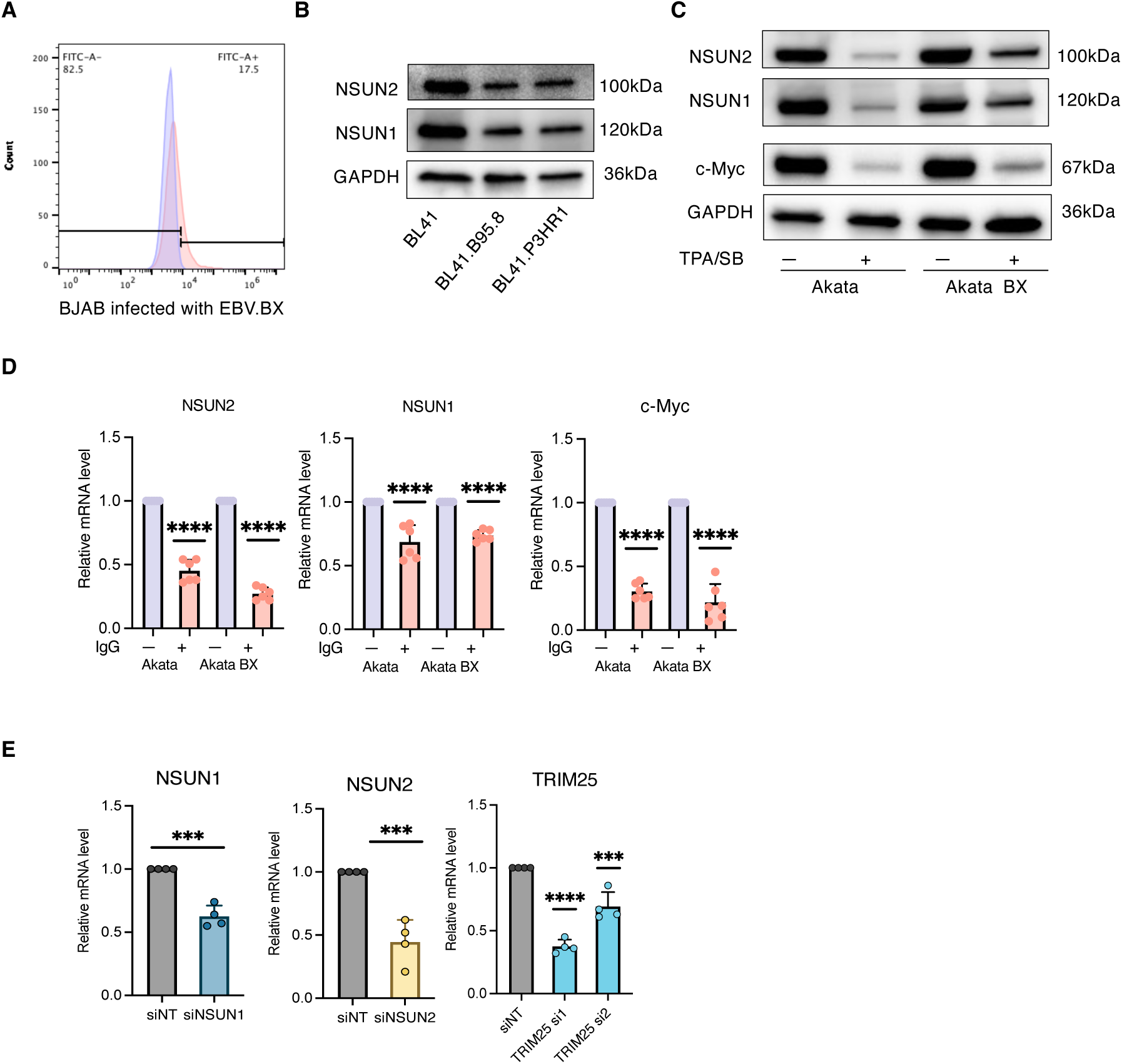
(A) BJAB cells were inoculated with EBV.BX viruses. Percentage of GFP-expressing cells indicating EBV infection was assessed by flow cytometry analysis. (B) BL41.B958 and BL41.P3HR1 (EBV-positive) cells as well as their parental BL41 (EBV-naïve) cells were subjected to protein immunoblotting analysis of NSUN2/1 using their specific antibodies. GAPDH was a loading control. (C) Akata (EBV-naïve) and Akata BX (EBV-positive) cells were treated with TPA/NaB or mock, followed by protein immunoblotting analysis of NSUN2/1 and c-Myc. (D) Akata (EBV-naïve) and Akata BX (EBV-positive) cells were treated with human IgG or mock, followed by RT-qPCR analysis of NSUN2/1 and c-Myc, which was normalized to GAPDH. (E) Akata cells were transiently transfected with siRNAs targeting NSUN2/1or TRIM25, or siNT, followed by RT-qPCR analysis of their expression to confirm knockdown efficiency. qPCR results from at least two independent experiments were presented as mean ± SD. (\**p* < 0.05, \*\**p* < 0.01, \*\*\**p* < 0.001, \*\*\*\**p* < 0.0001, Student’s t test).

**Figure S7.**
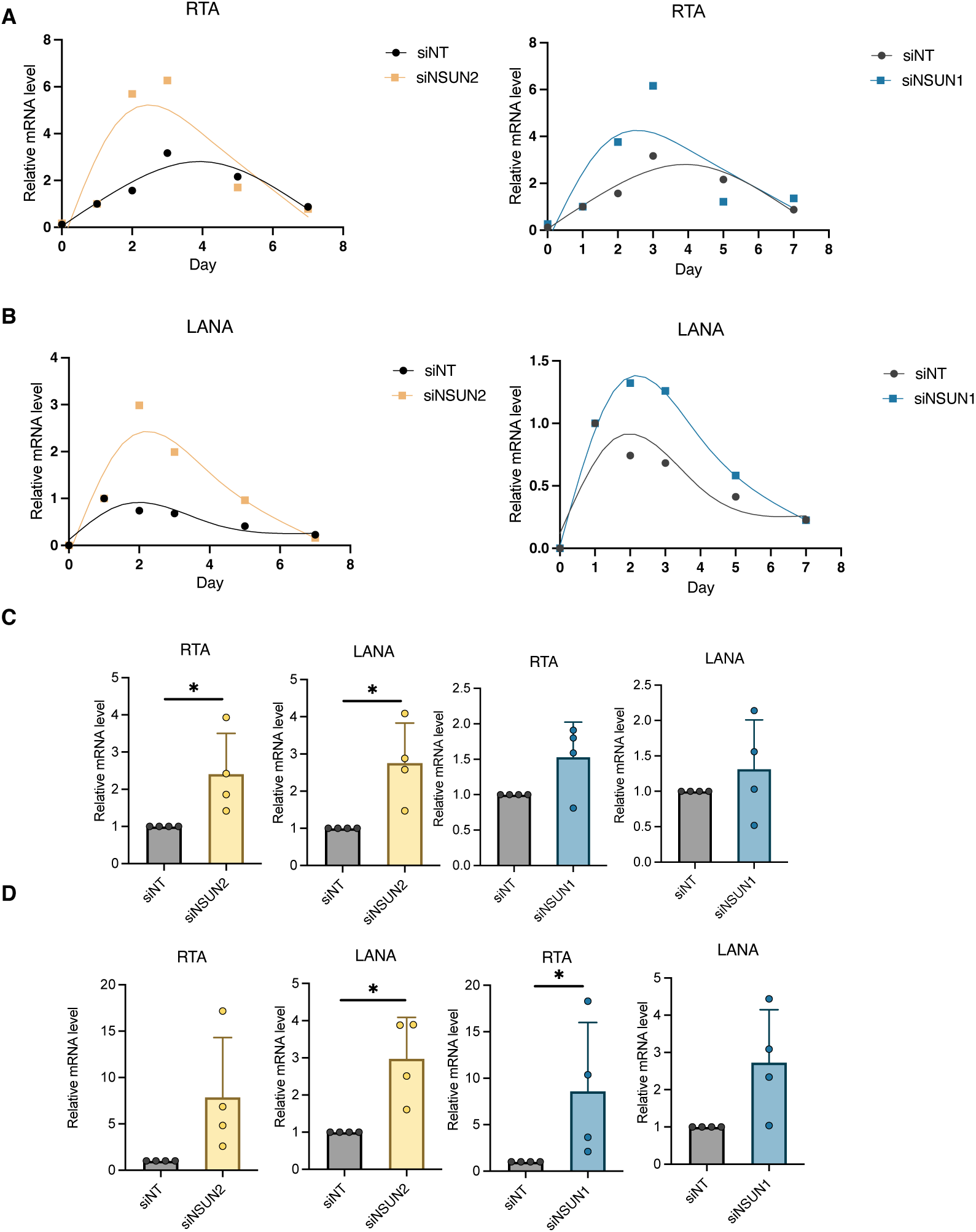
(A, B) TIME cells were transiently transfected with siRNAs targeting NSUN2/1, and inoculated with KSHV.BAC16 viruses. The cell lysates were harvested at indicated time points and subjected to RT-qPCR analysis to quantify KSHV RTA and LANA gene expression. (C) TREx.BCBL1.Rta cells were transiently transfected with siRNAs targeting NSUN2/1 by electroporation. The supernatants were collected at day 3 post electroporation, filtered by 0.22 uM filter, and titrated using TIME cells. KSHV-titrated TIME cells were collected at day 2 post infection and subjected to RT-qPCR analysis to quantify KSHV RTA and LANA gene expression. (D) TIME cells were transiently transfected with siRNAs targeting NSUN2/1 using RNAiMAX and inoculated with KSHV.BAC16 viruses. The supernatants were collected at day 2 post infection, filtered by 0.22 uM filter, and titrated using the fresh batch of TIME cells. KSHV-titrated TIME cells were collected at day 2 post infection and subjected to RT-qPCR analysis to quantify KSHV RTA and LANA gene expression. qPCR results from at least two independent experiments were presented as mean ± SD. (\**p* < 0.05, \*\**p* < 0.01, \*\*\**p* < 0.001, \*\*\*\**p* < 0.0001, Student’s t test).

**Figure S8.**
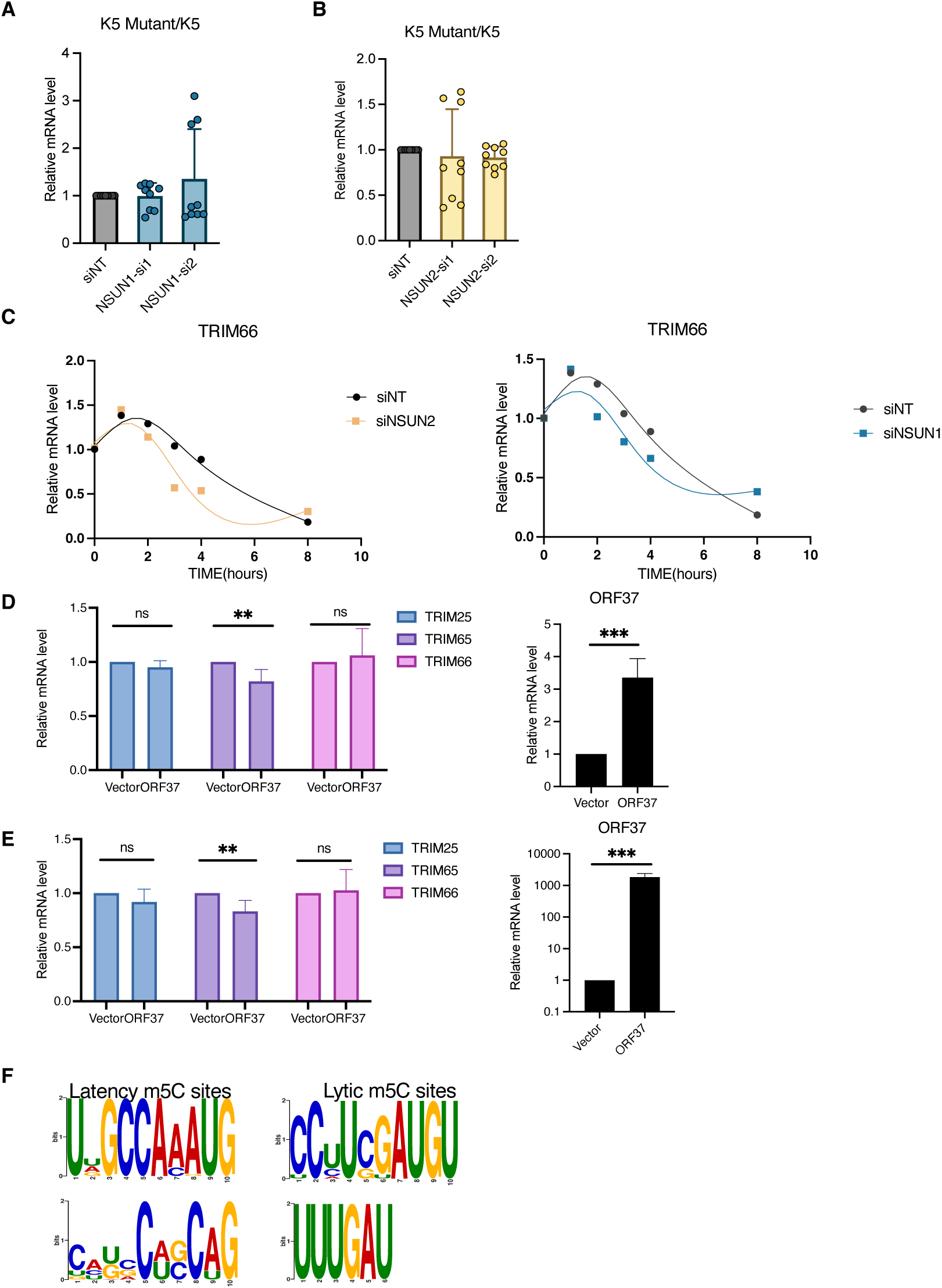
(A, B) HEK293T cells were transiently transfected with siRNAs targeting NSUN2/1, and transfected with the pcDNA3.1 vector expressing either wild-type or 544CtoA mutated KSHV K5 cDNA. The cell lysates were collected and subjected to RT-qPCR analysis of K5 mRNA level. The ratio of mRNA level for K5 wild-type *vs* 544CtoA mutant was calculated. (C) TREx.BCBL1.Rta cells were transiently transfected with siRNAs targeting NSUN2/1 by electroporation, followed by Actinomycin D treatment. Cells were collected at the indicated time points. Total RNAs were extracted and subjected to RT-qPCR analysis of TRIM66 mRNA level. (D) TREx.BCBL1.Rta cells were transiently transfected with the pcDNA3.1 vector expressing KSHV SOX/ORF37 cDNA or the empty vector by electroporation. The cells were collected at day 3 post electroporation. Total RNAs were extracted and subjected to RT-qPCR analysis to quantify gene expression for TRIM25, TRIM65, TRIM66, or KSHV ORF37. (E) iSLK.BAC16 cells were transiently transfected with the pcDNA3.1 vector expressing KSHV SOX/ORF37 or empty vector by Fugene 6. The cells were collected at day 2 post transfection. Total RNAs were extracted and subjected to RT-qPCR analysis to quantify gene expression for TRIM25, TRIM65, TRIM66, or KSHV ORF37. (F) Flanking sequencing of m5C sites identified from the condition of KSHV latency or KSHV lytic reactivation were used as input for *de novo* RNA motif finding by using MEME program. qPCR results from at least two independent experiments were presented as mean ± SD. (\**p* < 0.05, \*\**p* < 0.01, \*\*\**p* < 0.001, \*\*\*\**p* < 0.0001, Student’s t test).

**Figure S9.**
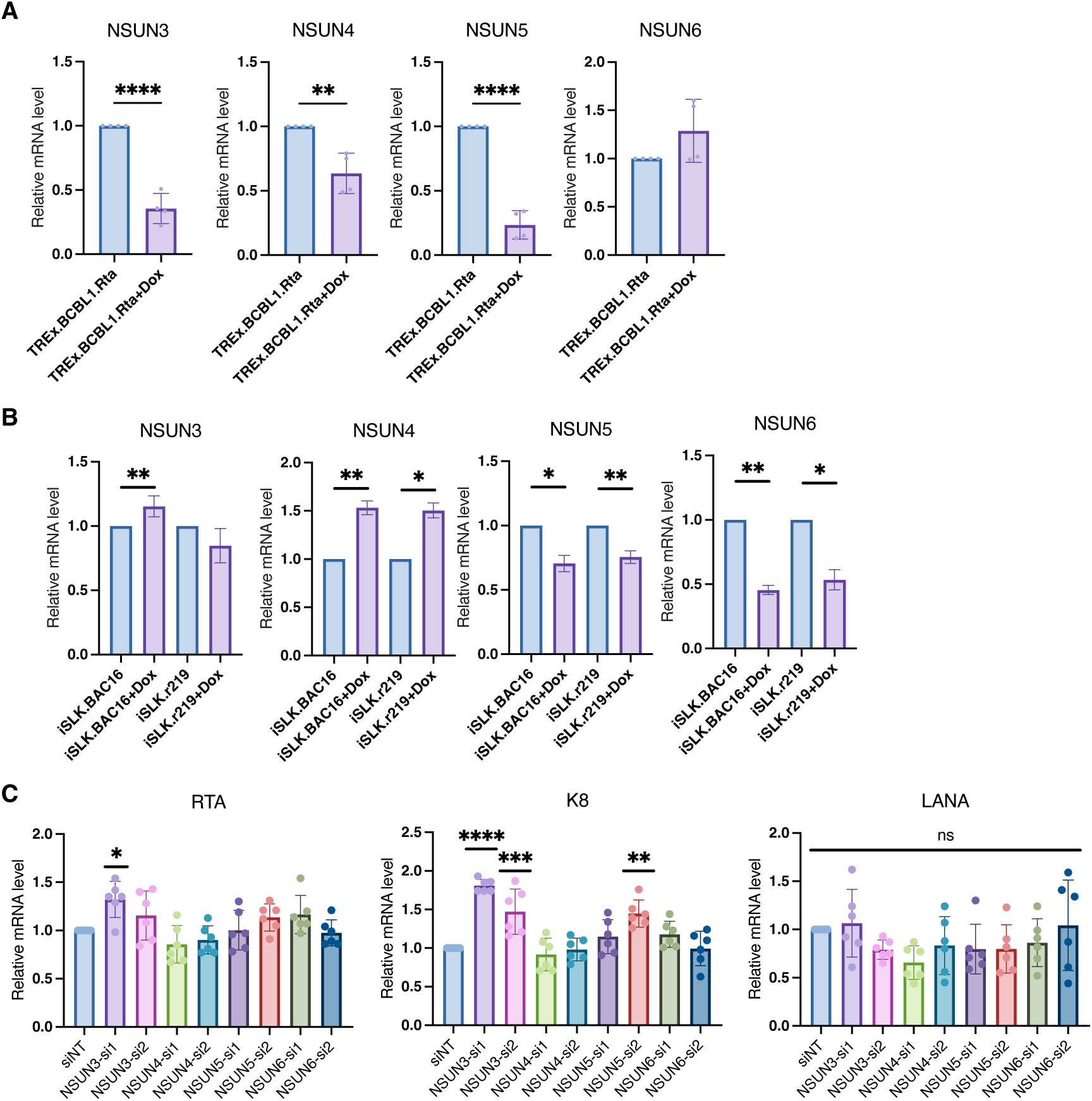
(A, B) TREx.BCBL1.Rta cells (A), iSLK.BAC16 and iSLK.r219 cells (B) were treated with Doxycycline or mock. The RNAs extracted from above cells were subjected to RT-qPCR analysis for quantification of NSUN3-6 gene expression, which was normalized to GAPDH. (C) TREx.BCBL1.Rta cells were transiently transfected with siRNAs targeting NSUN3-6 by electroporation. Total RNAs were extracted and subjected to RT-qPCR analysis of KSHV RTA, K8, LANA gene expression. qPCR results from at least two independent experiments were presented as mean ± SD. (\**p* < 0.05, \*\**p* < 0.01, \*\*\**p* < 0.001, \*\*\*\**p* < 0.0001, Student’s t test).

**Figure S10.**
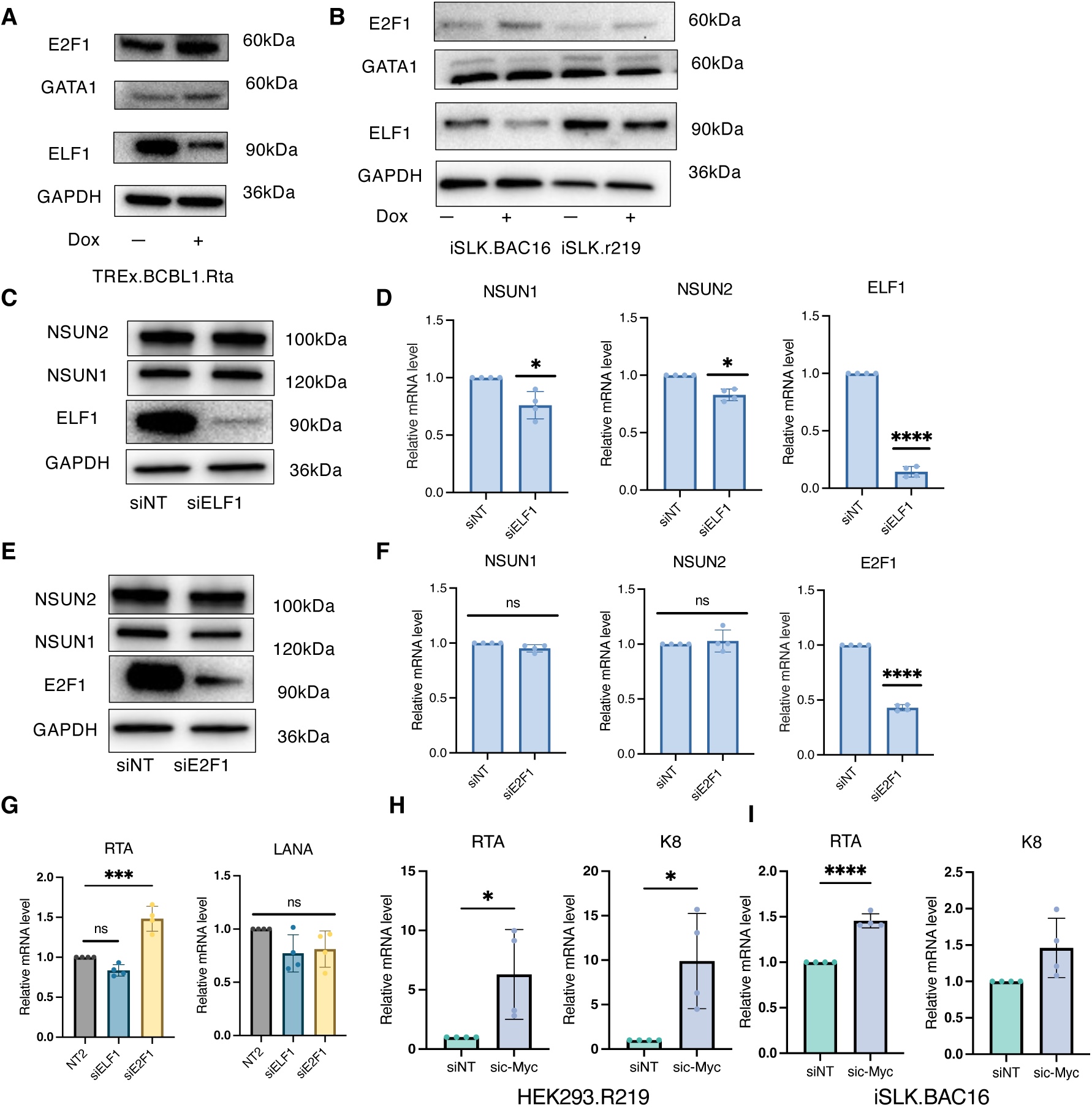
(A, B) TREx.BCBL1.Rta cells (A), iSLK.BAC16 and iSLK.r219 cells (B) were treated with Doxycycline or mock. The lysates of above cells were subjected to protein immunoblotting analysis by using the antibodies recognizing GATA1, ELF1, E2F1. GAPDH was used as a loading control. (C-F) TREx.BCBL1.Rta cells were transiently transfected with siRNAs targeting ELF1 or E2F1 by electroporation. The lysates of above samples were subjected to protein immunoblotting analysis by using the antibodies recognizing NSUN1/2, E2F1, or ELF1 (C, E). GAPDH was used as a loading control. Total RNAs were extracted from the above cells as well and subjected to RT-qPCR analysis of NSUN1/2, ELF1, E2F1 gene expression (D, F). (G) TREx.BCBL1.Rta cells were transiently transfected with siRNAs targeting ELF1 or E2F1 by electroporation. Total RNAs were extracted and subjected to RT-qPCR analysis of KSHV RTA and LANA gene expression. (H, I) HEK293.R219 (H) or iSLK.BAC16 (I) cells were transiently transfected with siRNAs targeting c-Myc. Total RNAs were extracted and subjected to RT-qPCR analysis of KSHV RTA, K8, and LANA gene expression. qPCR results from at least two independent experiments were presented as mean ± SD. (\**p* < 0.05, \*\**p* < 0.01, \*\*\**p* < 0.001, \*\*\*\**p* < 0.0001, Student’s t test).

**Table S1.** Public-domain RNA-seq datasets used for the reanalysis.

| GEO Access | Cell line | Treatment |
| --- | --- | --- |
| GSE123898 | Trex.BCBL1.3XFLAG.RTA | Dox |
| GSE212350 | TREx-BCBL1-PAN-GFP | Dox |
| GSE172275 | iSLK.Bac16 | Dox & NaB |
| GSE200781 | iSLK.R219 | Dox |
| GSE201000 | iSLK.Bac16 | Dox |
| GSE194239 | BCBL1 | VA |

Public RNA-seq for EBV reactivation
| GEO Access | Cell line | Treatment |
| --- | --- | --- |
| GSE237484 | Lymphoblastoid cell line | TPA & NaB |
| GSE240008 | Mutu | IgM |
| GSE240008 | Mutu | ZTA over-expression |
| GSE240008 | AKATA | IgG |

**Table S2.**
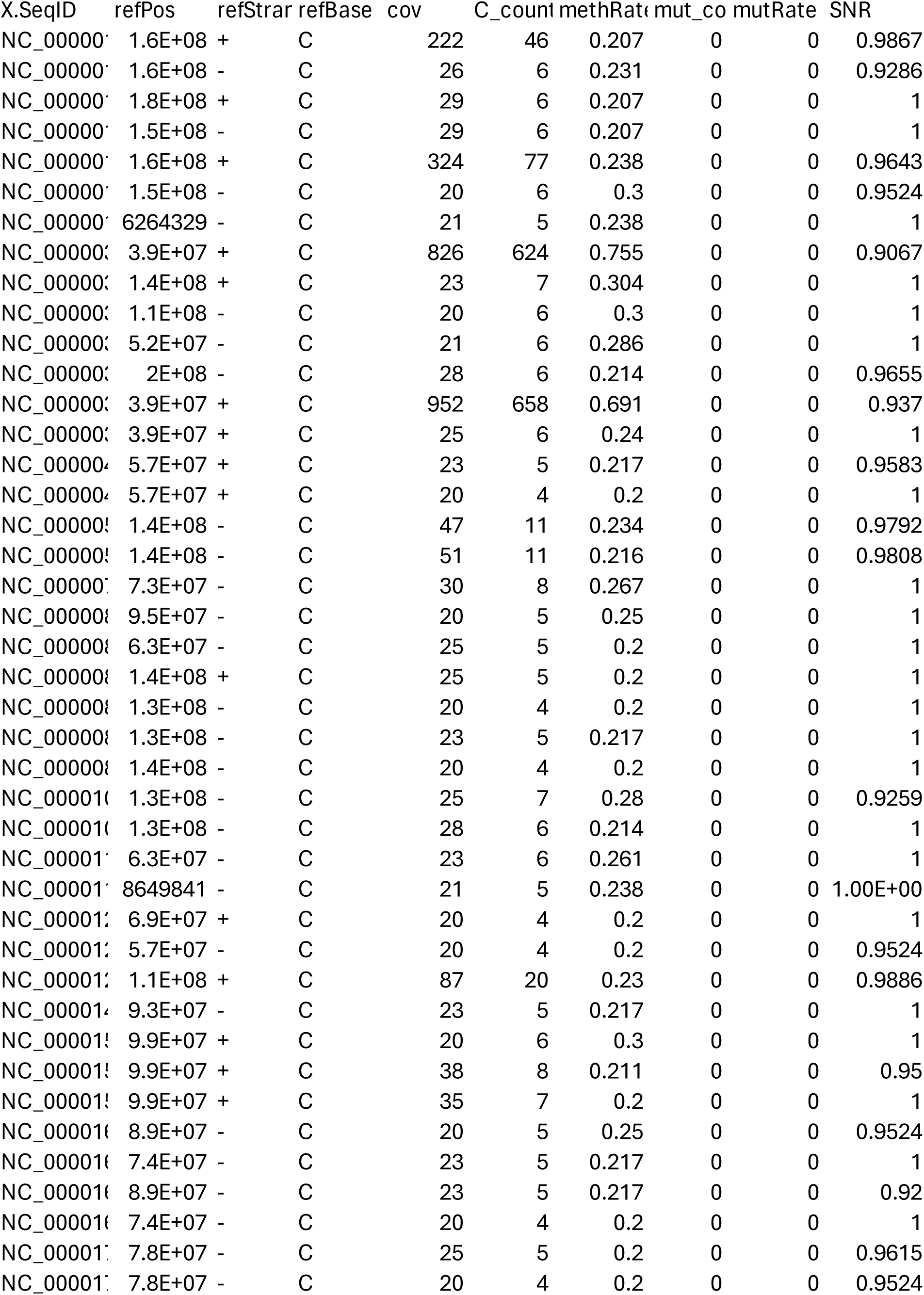
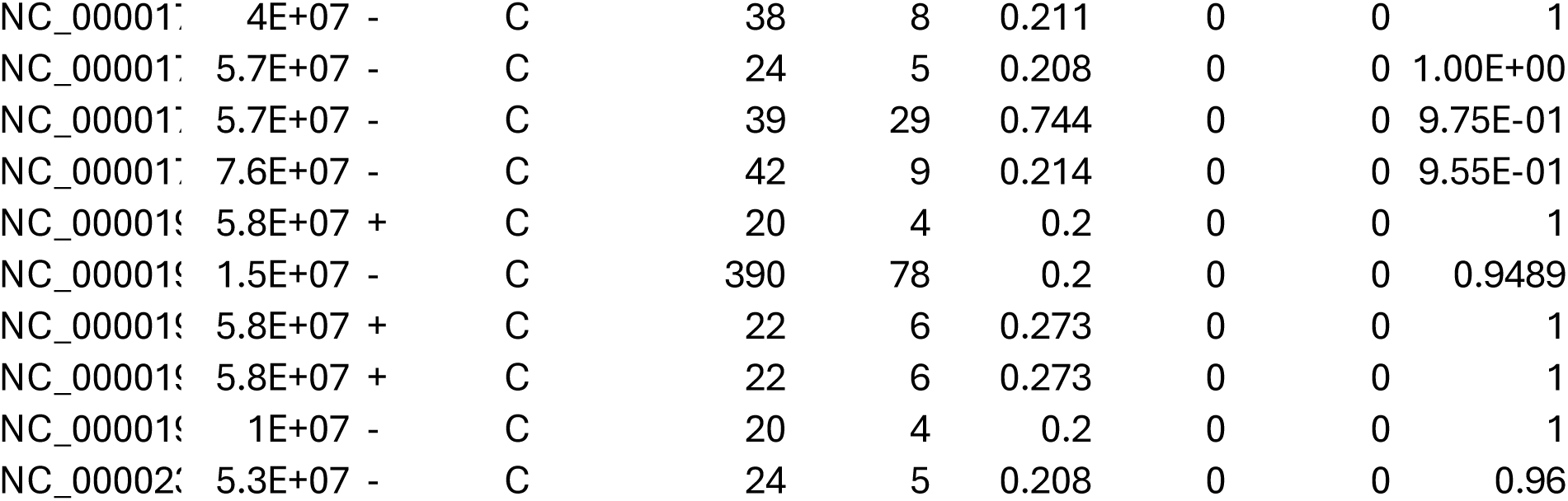
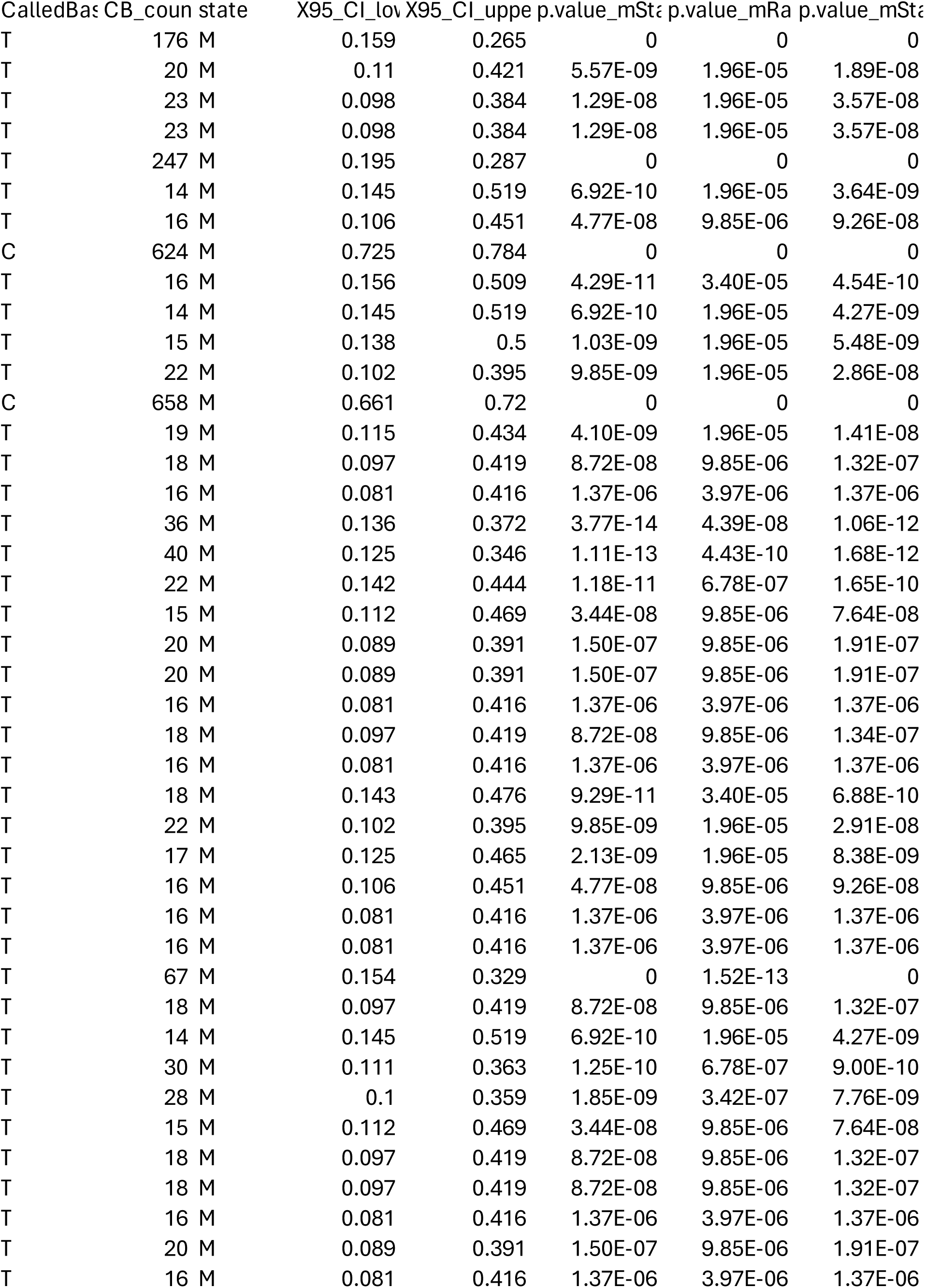
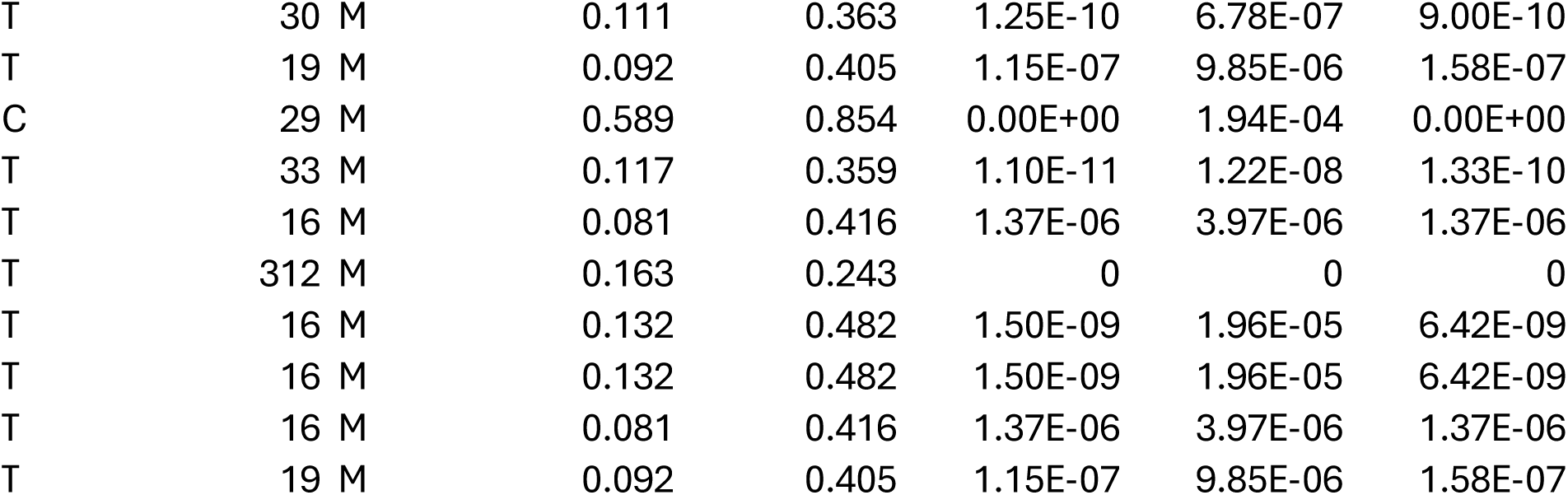
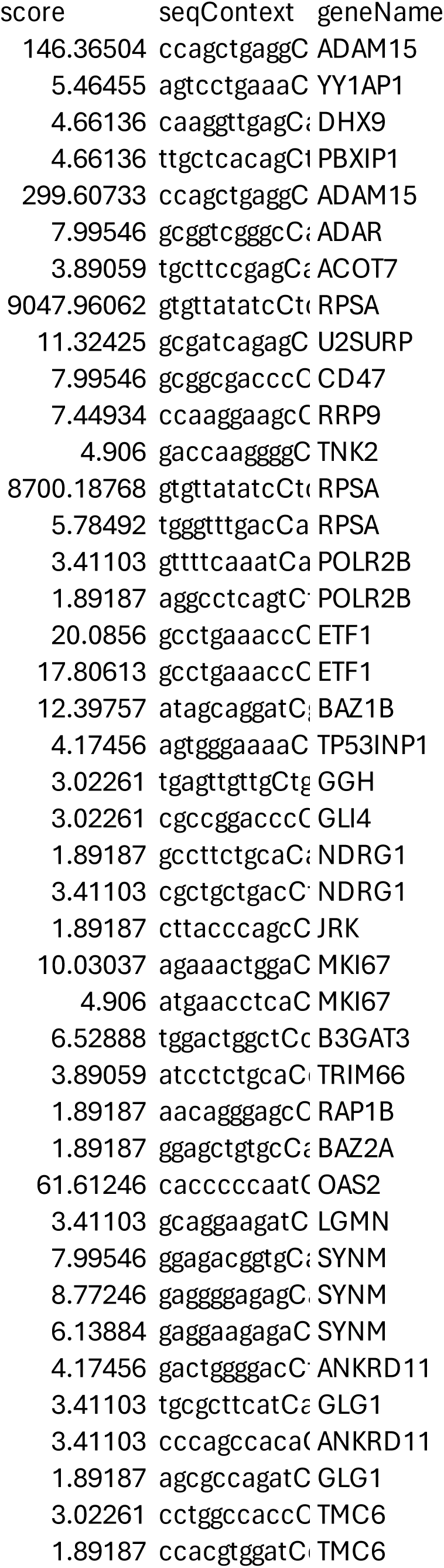
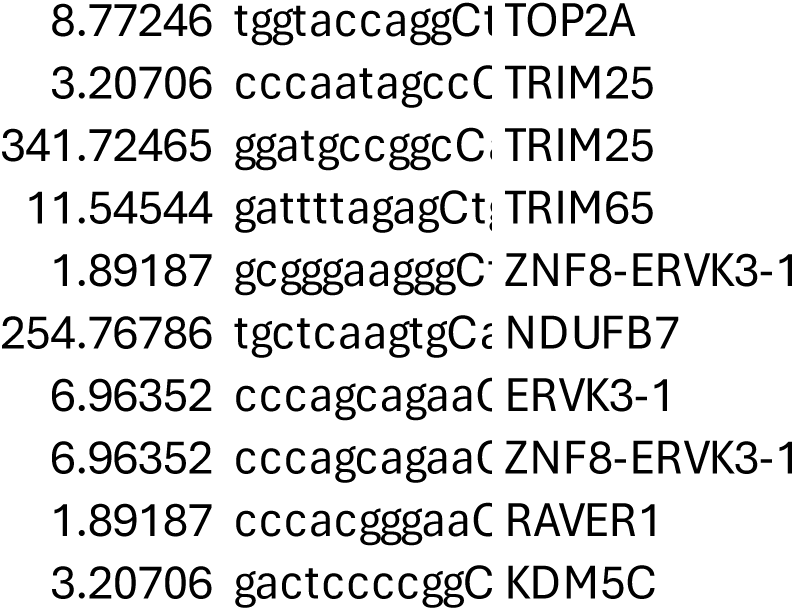
Identified m5C sites of host mRNAs subjected to the regulation of KSHV infection status.

**Table S3.**
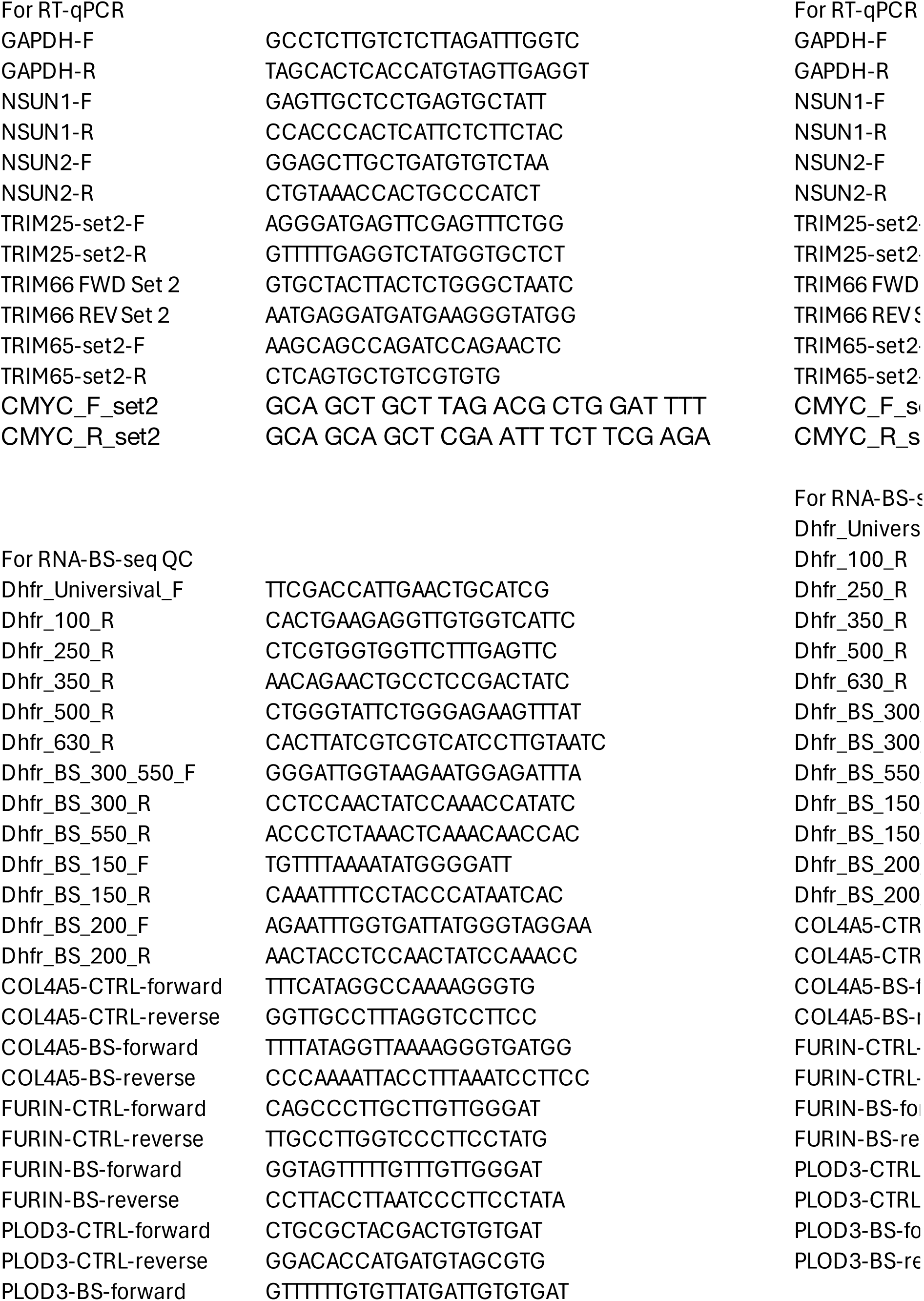
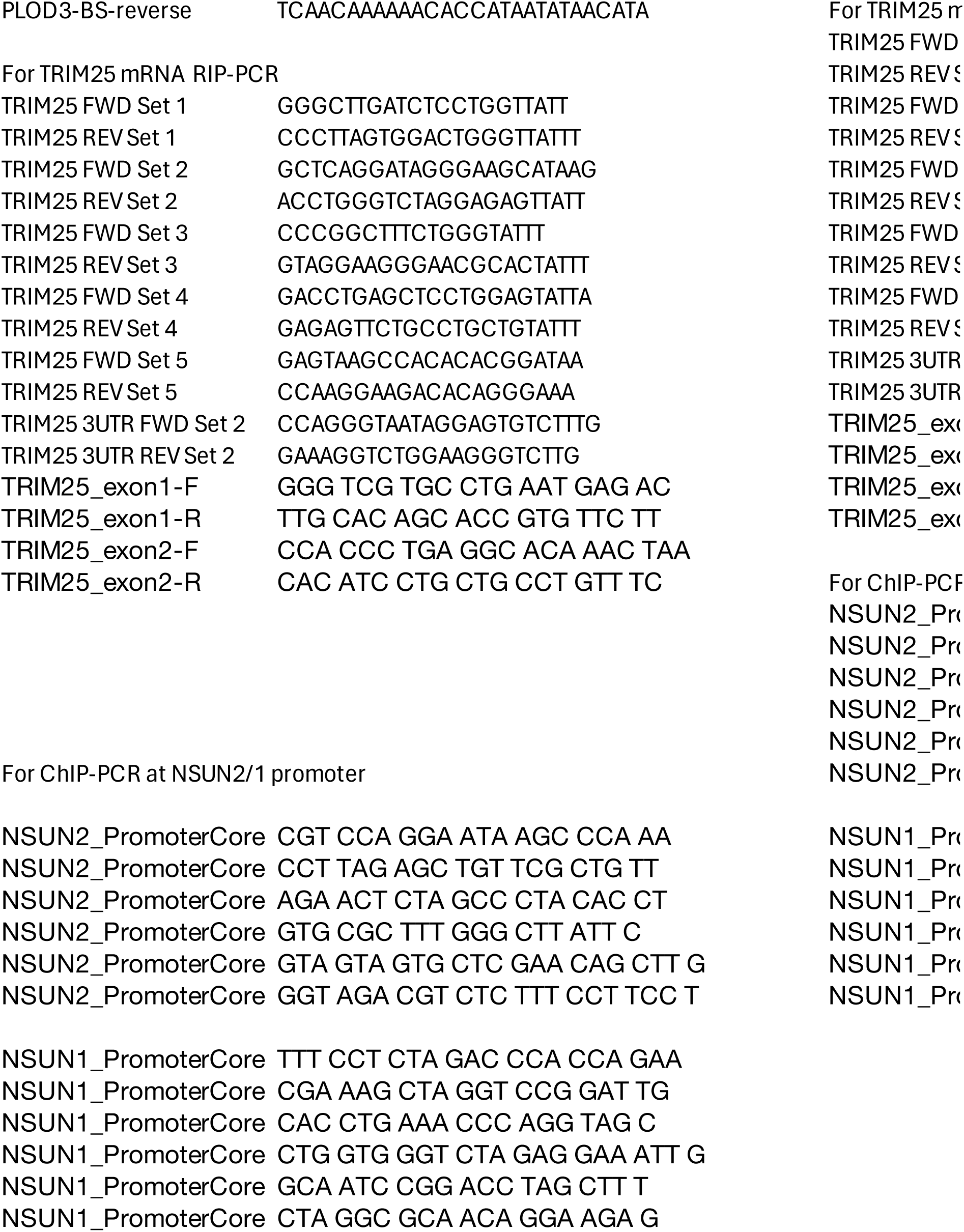
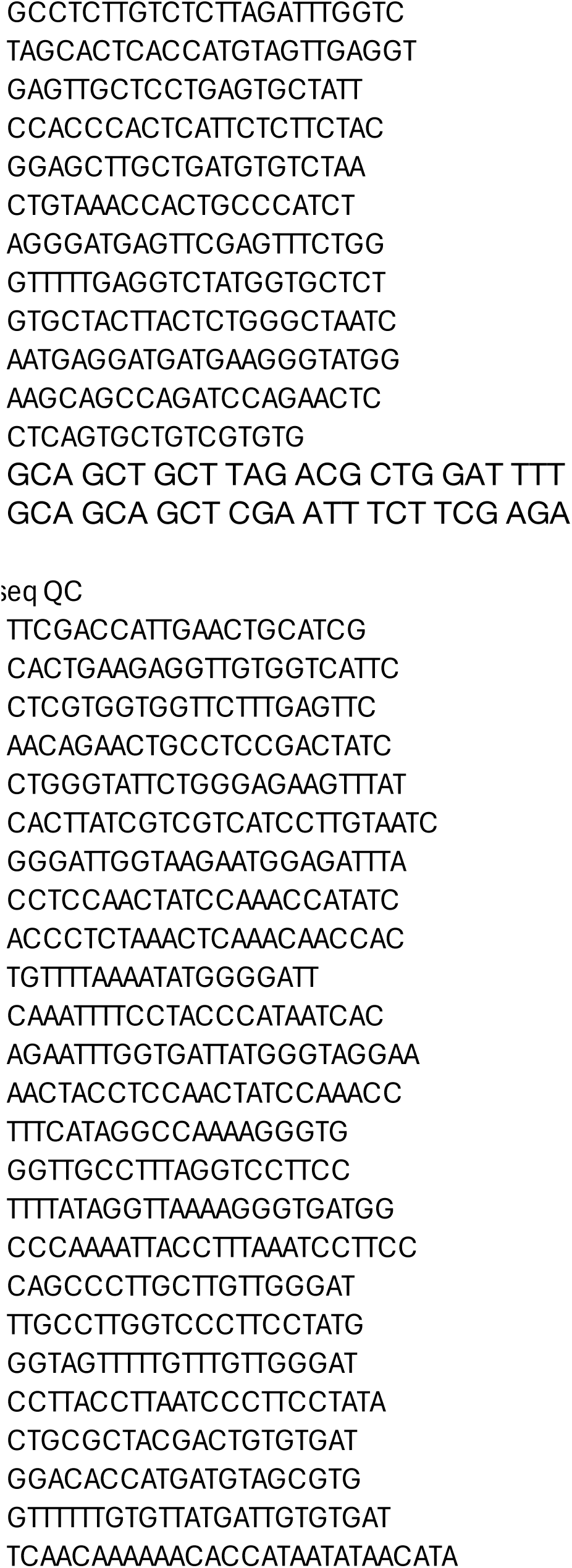
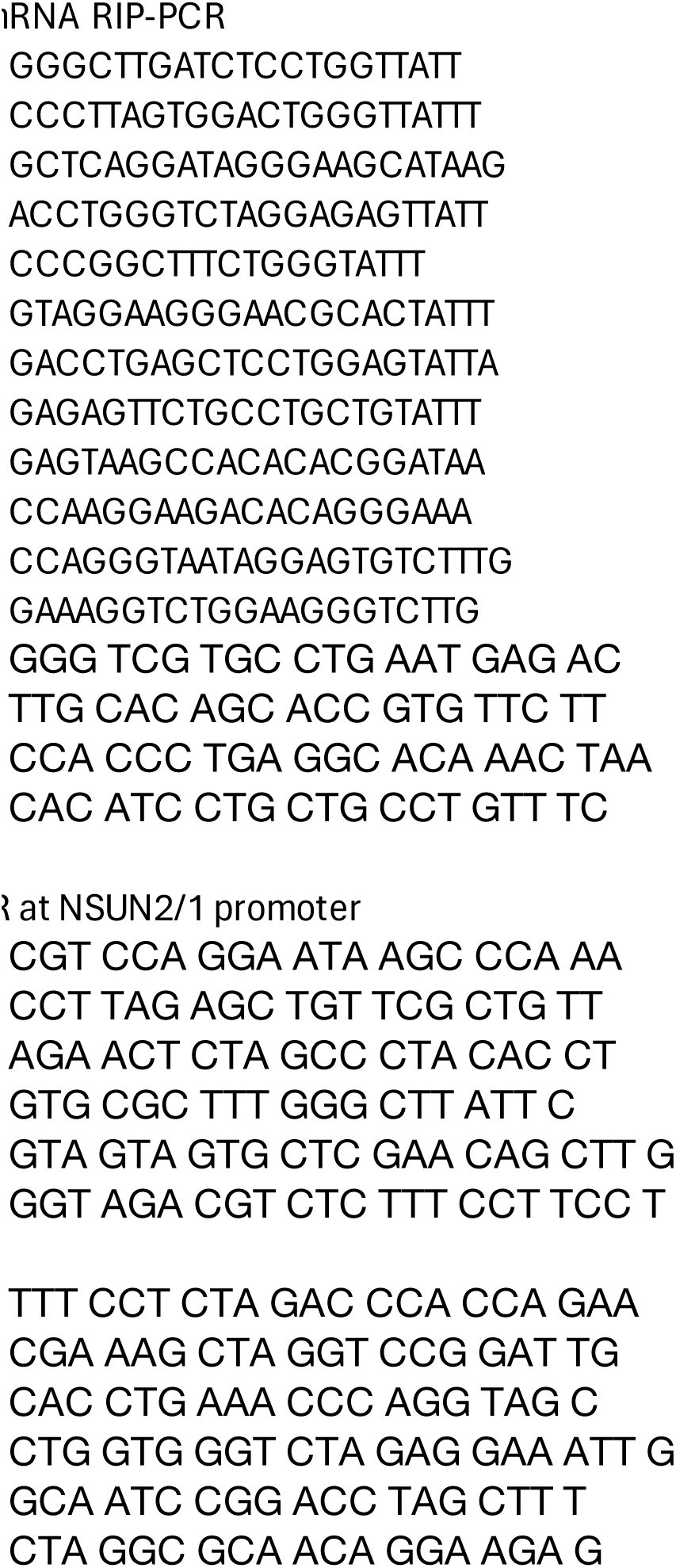
Sequence of primers used in the study.

**Table S4.**
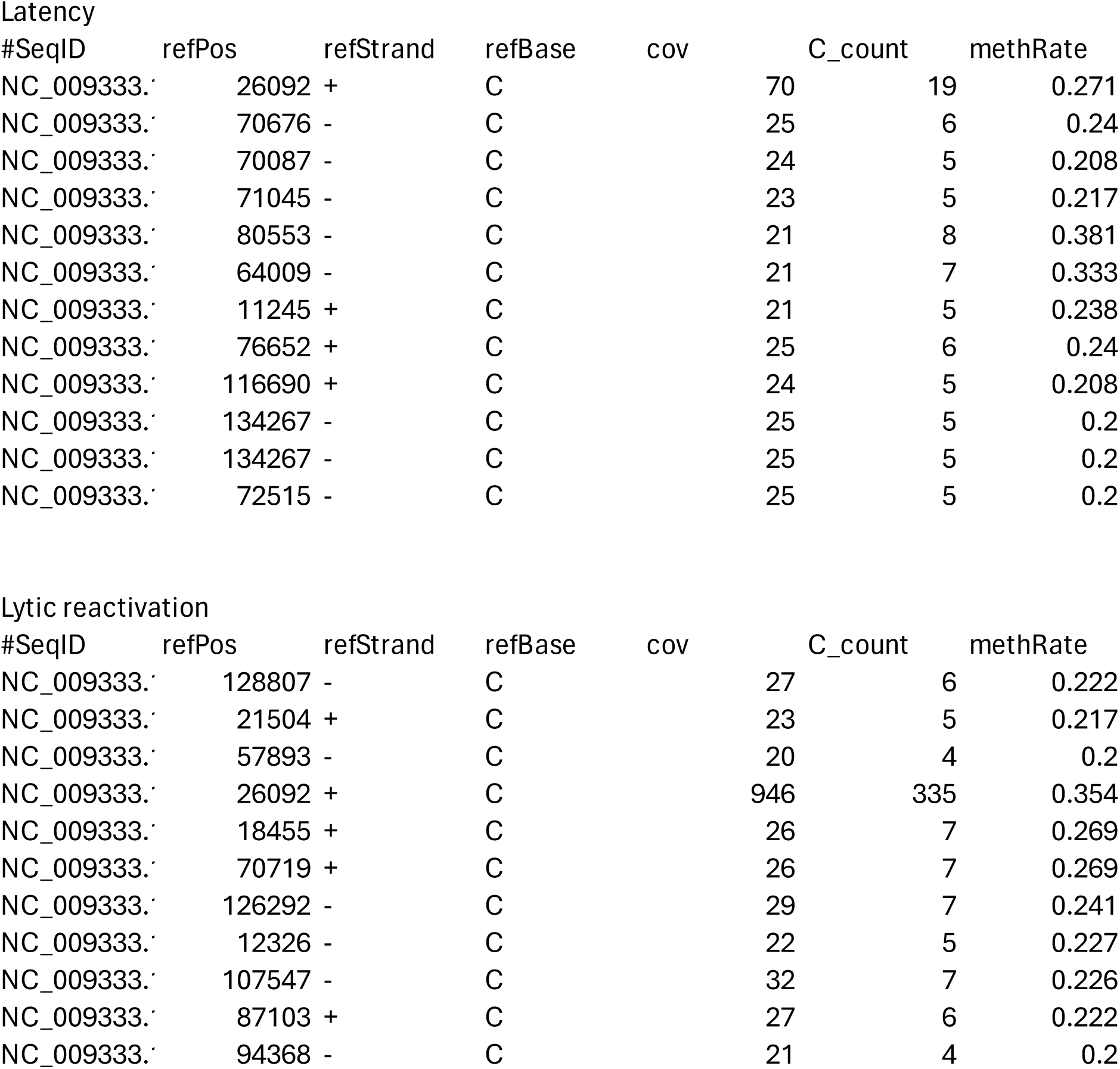
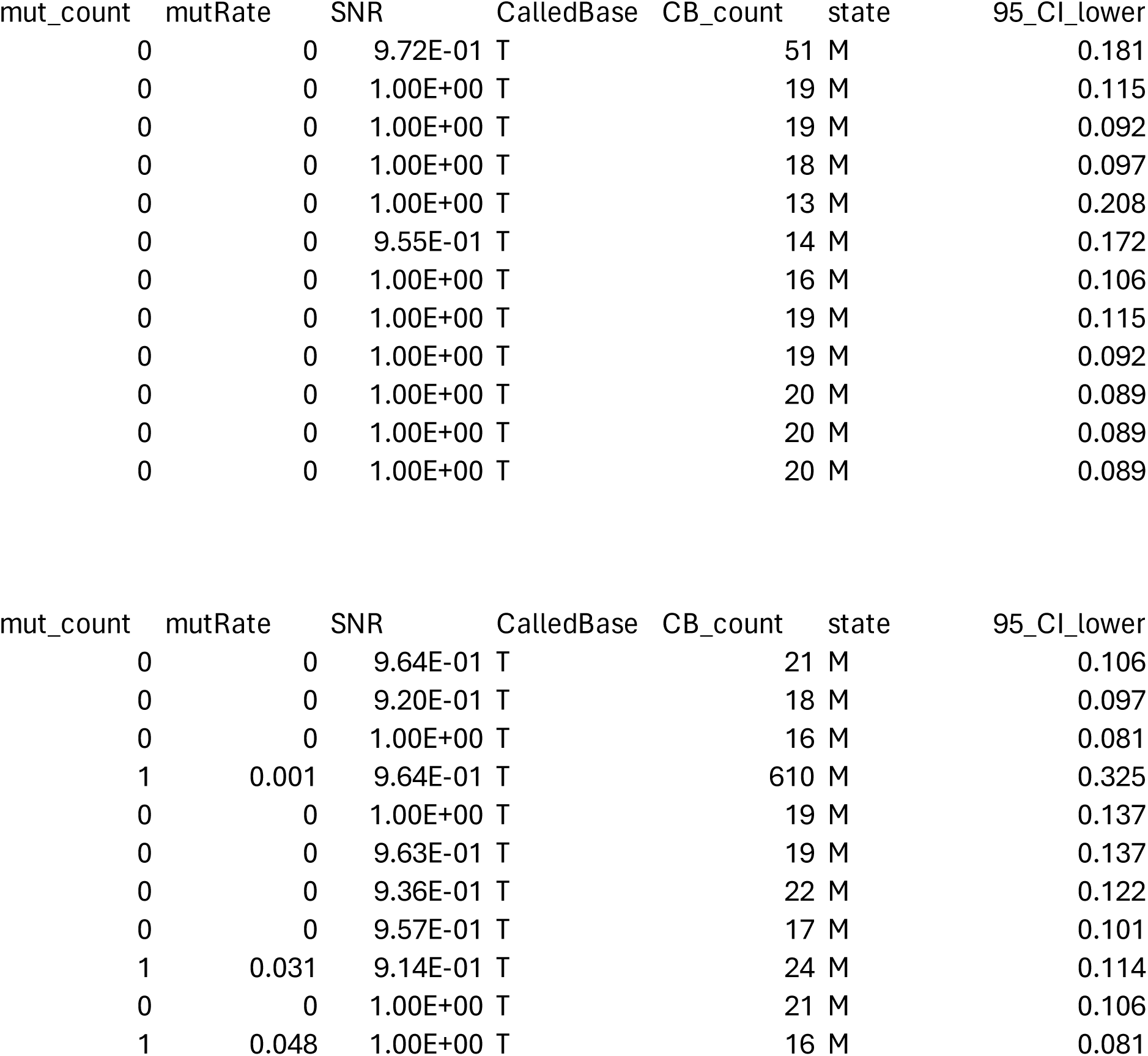
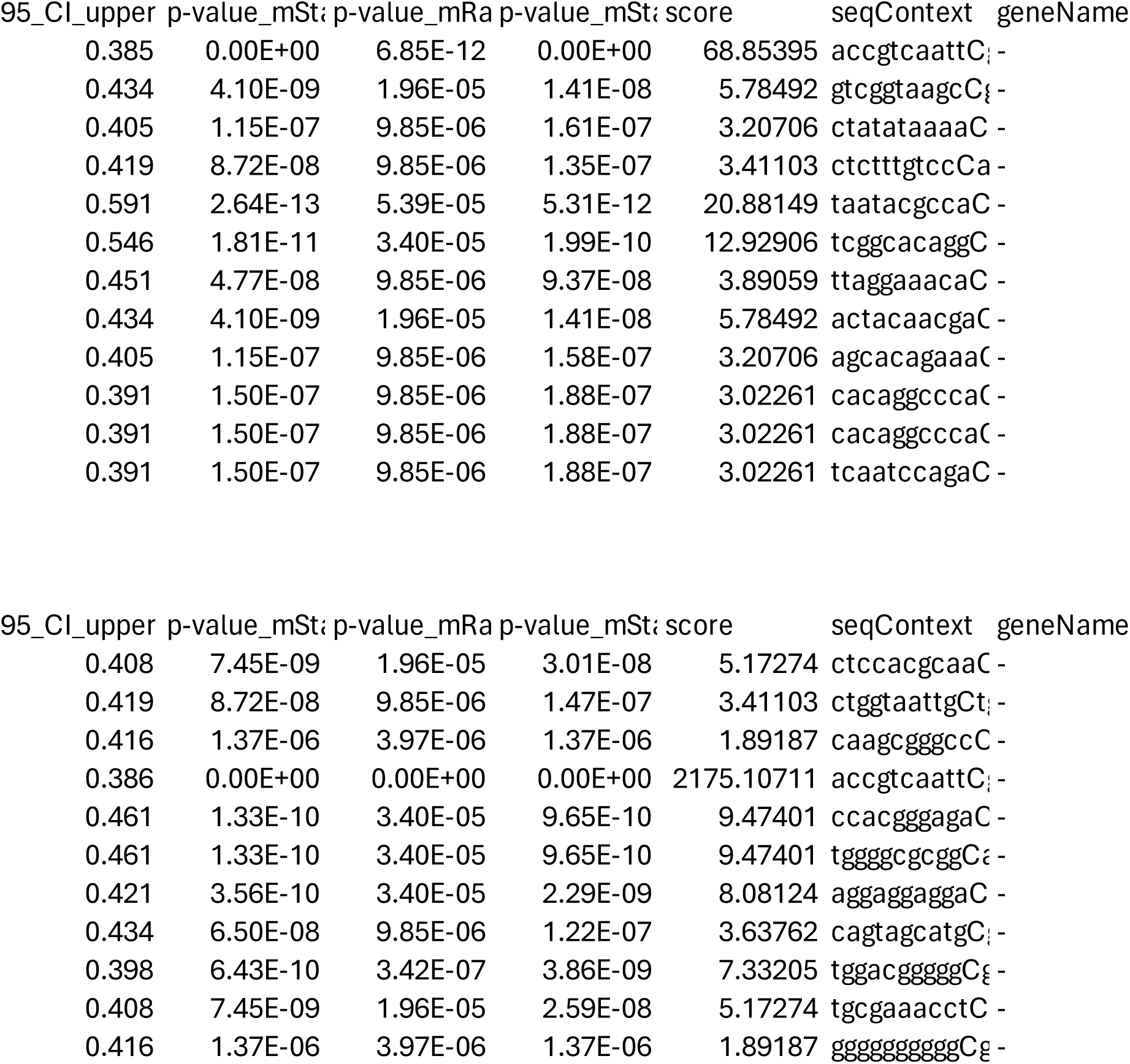
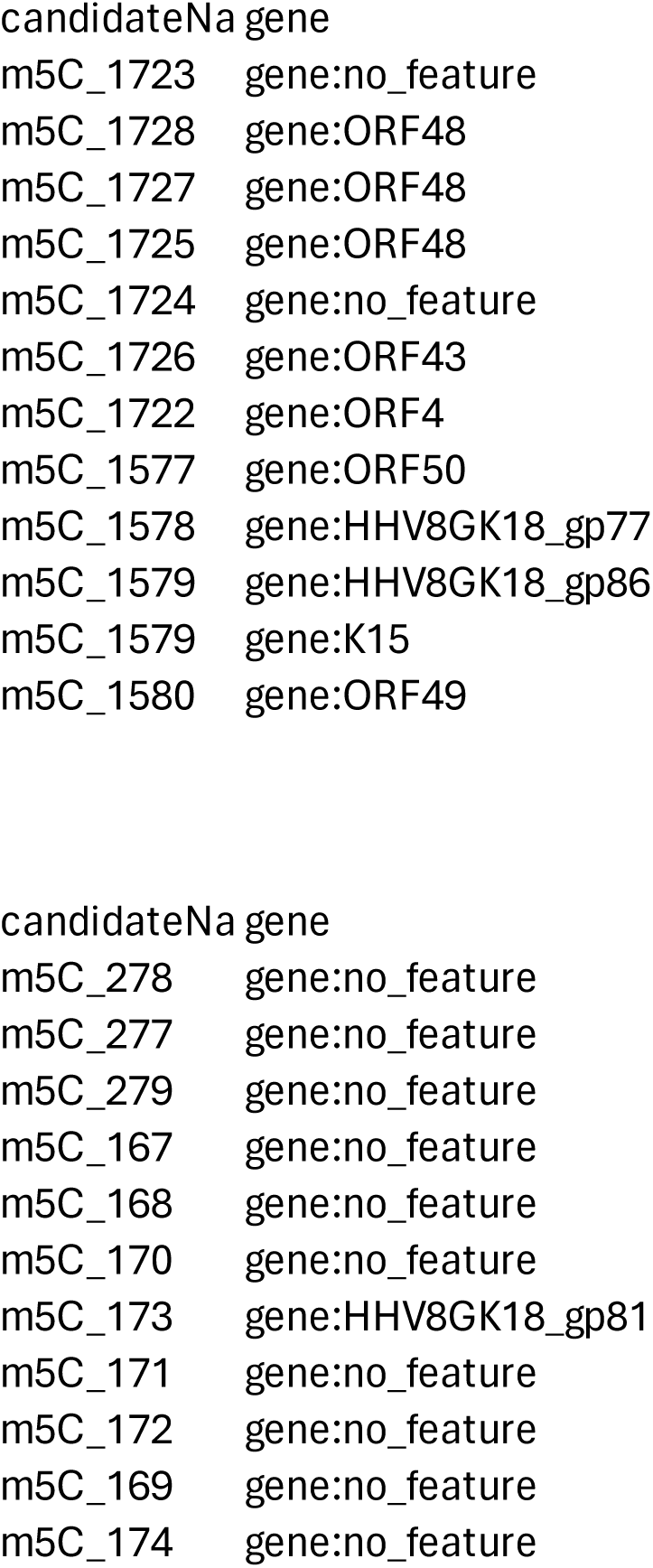
Identified m5C sites of viral mRNAs that occur during KSHV latency and lytic reactivation.

**Table S5.** Expression of RNA m5C writers during KSHV latency vs lytic reactivation by reanalyzing public-domain RNA-seq datasets.

| Symbol | Fold change | padj | Dataset |
| --- | --- | --- | --- |
| NSUN3 | 0.9369032 | 0.418 | GSE123898 |
| NSUN3 | 0.68152052 | 0.186 | GSE172275 |
| NSUN3 | 1.07118444 | 0.324 | GSE194239 |
| NSUN3 | 0.72162089 | 0.0961 | GSE200781 |
| NSUN3 | 1.16315737 | 0.212 | GSE212350 |
| NSUN3 | 0.68074454 | 0.20823131 | GSE201000 |
| NSUN4 | 1.16548586 | 0.000227 | GSE123898 |
| NSUN4 | 0.73847296 | 0.125 | GSE172275 |
| NSUN4 | 0.73383756 | 2.97E-09 | GSE194239 |
| NSUN4 | 0.96459551 | 0.841 | GSE200781 |
| NSUN4 | 0.98567839 | 0.897 | GSE212350 |
| NSUN4 | 0.91492286 | 0.63254663 | GSE201000 |
| NSUN5 | 0.87097125 | 4.80E-05 | GSE123898 |
| NSUN5 | 0.48908903 | 1.29E-05 | GSE172275 |
| NSUN5 | 0.72733408 | 1.59E-12 | GSE194239 |
| NSUN5 | 0.55847786 | 1.51E-05 | GSE200781 |
| NSUN5 | 0.60965607 | 1.80E-16 | GSE212350 |
| NSUN5 | 0.69014089 | 0.04961975 | GSE201000 |
| NSUN6 | 0.96144191 | 0.5 | GSE123898 |
| NSUN6 | 1.24340217 | 0.587 | GSE172275 |
| NSUN6 | 1.12492009 | 0.0671 | GSE194239 |
| NSUN6 | 0.92625671 | 0.807 | GSE200781 |
| NSUN6 | 1.27754419 | 0.00285 | GSE212350 |
| NSUN6 | 0.66491442 | 0.18657781 | GSE201000 |
| NSUN7 | 0.86340072 | 0.641 | GSE123898 |
| NSUN7 | 2.57092585 | 2.11E-07 | GSE172275 |
| NSUN7 | 2.82001727 | 1.96E-12 | GSE200781 |
| NSUN7 | 0.82377022 | 0.87963447 | GSE201000 |
| DNMT1 | 0.88536301 | 1.43E-08 | GSE123898 |
| DNMT1 | 0.85624526 | 0.337 | GSE172275 |
| DNMT1 | 0.61026949 | 8.71E-43 | GSE194239 |
| DNMT1 | 0.40710984 | 3.09E-13 | GSE200781 |
| DNMT1 | 0.77393242 | 1.42E-10 | GSE212350 |
| DNMT1 | 1.16997441 | 0.42099578 | GSE201000 |
| TRDMT1 (DN | 0.70281705 | 1.61E-07 | GSE123898 |
| TRDMT1 | 0.98597844 | 0.973 | GSE172275 |
| TRDMT1 | 1.80345522 | 2.41E-15 | GSE194239 |
| TRDMT1 | 1.01825295 | 0.949 | GSE200781 |
| TRDMT1 | 0.80066103 | 0.056 | GSE212350 |
| TRDMT1 | 0.67025736 | 0.40742946 | GSE201000 |
| DNMT3A | 0.66556243 | 2.89E-12 | GSE123898 |
| DNMT3A | 1.01866381 | 0.958 | GSE172275 |
| DNMT3A | 1.13541206 | 0.021 | GSE194239 |
| DNMT3A | 1.17973023 | 0.378 | GSE200781 |
| DNMT3A | 1.09976746 | 0.296 | GSE212350 |
| DNMT3A | 0.99832242 | 0.99462494 | GSE201000 |
| DNMT3B | 0.85281197 | 0.132 | GSE123898 |
| DNMT3B | 3.11851241 | 1.51E-08 | GSE172275 |
| DNMT3B | 1.3433996 | 0.00167 | GSE194239 |
| DNMT3B | 0.50348545 | 5.88E-05 | GSE200781 |
| DNMT3B | 0.66542893 | 0.0157 | GSE212350 |
| DNMT3B | 2.65024359 | 6.98E-16 | GSE201000 |
| NSUN1 | 0.85559959 | 2.77E-09 | GSE123898 |
| NSUN1 | 0.45860246 | 5.49E-09 | GSE172275 |
| NSUN1 | 0.77586881 | 1.23E-12 | GSE194239 |
| NSUN1 | 0.4779127 | 6.36E-10 | GSE200781 |
| NSUN1 | 0.7476211 | 1.44E-09 | GSE212350 |
| NSUN1 | 0.84425157 | 0.00622153 | GSE201000 |
| NSUN2 | 0.88700196 | 2.71E-05 | GSE123898 |
| NSUN2 | 0.58406992 | 0.00528 | GSE172275 |
| NSUN2 | 0.8540151 | 9.05E-06 | GSE194239 |
| NSUN2 | 0.55226553 | 1.72E-06 | GSE200781 |
| NSUN2 | 0.80932642 | 0.00024 | GSE212350 |
| NSUN2 | 0.65628712 | 0.00248943 | GSE201000 |

